# Pan-cancer proteomic map of 949 human cell lines reveals principles of cancer vulnerabilities

**DOI:** 10.1101/2022.02.26.482008

**Authors:** Emanuel Gonçalves, Rebecca C Poulos, Zhaoxiang Cai, Syd Barthorpe, Srikanth S Manda, Natasha Lucas, Alexandra Beck, Daniel Bucio-Noble, Michael Dausmann, Caitlin Hall, Michael Hecker, Jennifer Koh, Sadia Mahboob, Iman Mali, James Morris, Laura Richardson, Akila J Seneviratne, Erin Sykes, Frances Thomas, Sara Valentini, Steven G Williams, Yangxiu Wu, Dylan Xavier, Karen L MacKenzie, Peter G Hains, Brett Tully, Phillip J Robinson, Qing Zhong, Mathew J Garnett, Roger R Reddel

**Affiliations:** Wellcome Sanger Institute, Wellcome Genome Campus, Cambridge, CB10 1SA, UK; Instituto Superior Técnico (IST), Universidade de Lisboa, 1049-001, Lisboa, Portugal; INESC-ID, 1000-029, Lisboa, Portugal; ProCan^®^, Children’s Medical Research Institute, Faculty of Medicine and Health, The University of Sydney, Westmead, NSW, Australia

**Keywords:** Proteomics, Cancer, Cell line, Mass spectrometry, Drug response, CRISPR-Cas9, Gene essentiality, Cancer vulnerability

## Abstract

The proteome provides unique insights into biology and disease beyond the genome and transcriptome. Lack of large proteomic datasets has restricted identification of new cancer biomarkers. Here, proteomes of 949 cancer cell lines across 28 tissue types were analyzed by mass spectrometry. Deploying a clinically-relevant workflow to quantify 8,498 proteins, these data capture evidence of cell type and post-transcriptional modifications. Integrating multi-omics, drug response and CRISPR-Cas9 gene essentiality screens with a deep learning-based pipeline revealed thousands of protein-specific biomarkers of cancer vulnerabilities. Proteomic data had greater power to predict drug response than the equivalent portion of the transcriptome. Further, random downsampling to only 1,500 proteins had limited impact on predictive power, consistent with protein networks being highly connected and co-regulated. This pan-cancer proteomic map (ProCan-DepMapSanger), available at https://cellmodelpassports.sanger.ac.uk, is a comprehensive resource revealing principles of protein regulation with important implications for future clinical studies.

## Introduction

Precision medicine relies on the identification of specific molecular alterations that can stratify patients and guide the choice of effective therapeutic options. Cancer vulnerabilities, such as synthetic lethalities, can be systematically studied using functional genetic and small molecule screens. To circumvent the limitations of using patient tissue samples for this type of approach, biomarkers of cancer vulnerabilities have been analyzed using cancer cell lines, together with deep molecular characterization, functional genetic and pharmacological screens (Behan et al., 2019; Frejno et al., 2020; Ghandi et al., 2019; Iorio et al., 2016; Tsherniak et al., 2017). Direct measurement of proteins provides insights into the dynamic molecular behavior of cells, and can improve our understanding of genotype-to-phenotype relationships (Liu et al., 2016). Despite the development of precision oncology therapeutics, the complexity of cancer and the inability of genomics to accurately predict the proteome indicate that genomics alone is often insufficient to inform and guide the clinical care of many patients. Measurement of the proteome has the potential to expand understanding of cancer phenotypes, and to improve diagnosis and treatment choices.

Technological and methodological advances have enabled standardized quantification of thousands of proteins across dozens to hundreds of cell lines (Coscia et al., 2016; Gholami et al., 2013; Lawrence et al., 2015; Nusinow et al., 2020; Roumeliotis et al., 2017) and the profiling of clinical samples derived from minute tissue biopsies (Clark et al., 2019; Edwards et al., 2015; Frejno et al., 2017; Mertins et al., 2016; Pozniak et al., 2016; Tully, 2020; Vasaikar et al., 2019; Zhang et al., 2014, 2016). Using a data-independent acquisition (DIA)-mass spectrometry (MS) approach (Gillet et al., 2012; Guo et al., 2015; Lucas et al., 2019; Ludwig et al., 2018), together with a clinically-relevant sample processing workflow with novel data processing methods, it is now possible for proteomes to be acquired reproducibly at scale (Poulos et al., 2020; Tully et al., 2019). The generation and distribution of large-scale proteomic datasets have the potential to drive new computational approaches, including deep learning-based algorithms, to investigate the impact of molecular changes on cancer vulnerabilities. Such advancements will enable proteomics to contribute important clinical advances for cancer therapeutic applications.

Cell lines have been invaluable models for our understanding of cellular processes and the molecular drivers of carcinogenesis (Barretina et al., 2012; Behan et al., 2019; Garnett et al., 2012; Ghandi et al., 2019; Iorio et al., 2016; Meyers et al., 2017; Picco et al., 2019; Tsherniak et al., 2017), and for identifying cancer cell vulnerabilities to both genetic (Behan et al., 2019; Hart et al., 2015; McDonald et al., 2017; Meyers et al., 2017; Tsherniak et al., 2017) and pharmacological (Barretina et al., 2012; Corsello et al., 2020; Garnett et al., 2012; Iorio et al., 2016; Rees et al., 2016; Seashore-Ludlow et al., 2015) perturbations. However, proteomic quantifications for cancer cell lines are either limited in the range of cancer types or number of samples analyzed, or are largely unavailable (Frejno et al., 2020; Gholami et al., 2013; Guo et al., 2019; Li et al., 2017; Nusinow et al., 2020). For this reason, biomarkers for cancer vulnerabilities have so far been primarily based on genomic and transcriptomic measurements (Ghandi et al., 2019; Iorio et al., 2016; Tsherniak et al., 2017). Consequently, little is known about the contribution of the proteome to cancer vulnerabilities, or how the cancer proteome is regulated in diverse tissues and genetic contexts.

This study reports the largest pan-cancer cell line proteomic map compiled to date, quantifying 8,498 proteins across 949 cell lines. The generation and analysis of this rich resource involved the development of a clinically-oriented workflow with rapid sample processing and minimal complexity, followed by the application of a deep neural network-based computational pipeline to uncover novel cancer targets. Integration of our proteomic data (referred to as the ProCan-DepMapSanger dataset) with existing molecular and phenotypic datasets from the *Cancer Dependency Map* (Boehm et al., 2021), showed that protein networks are more strongly co-regulated than are transcriptomics and functional genomics. Our approach identified biomarkers of well-established cancer vulnerabilities and, more importantly, highlighted those that cannot be identified with genomic or transcriptomics alone. The proteome measured in our study had equivalent performance to the total measured transcriptome in predicting cancer phenotypes. Furthermore, random subsets of 1,500 proteins downsampled from the complete dataset achieved 88% of the power to predict drug responses. These results, and the clinical proteomic approach we present here, have broad implications for the design of future studies, ranging from basic research to clinical applications. Taken together, this study highlights the importance of including the proteome in multi-omic research, and its potentially pivotal role in providing novel treatments for cancer patients.

## Results

### A resource of 949 cancer cell line proteomes

To construct a pan-cancer proteomic map, proteomes of 949 human cancer cell lines were quantified from 28 tissues and over 40 genetically and histologically diverse cancer types (***Figure 1a*** and ***Figure S1a, Table S1a***). The proteome for each cell line was acquired by DIA-MS from six replicates using a clinically-relevant workflow that enables high throughput and minimal instrument downtime (see ***Methods, Figure S1b***). The resulting dataset was derived from 6,864 DIA-MS runs acquired over 10,000 MS hours (***Table S1b***), including peptide preparations derived from the human embryonic kidney cell line HEK293T that were used throughout all data acquisition periods and instruments for quality control. These data were deposited in the Proteomics Identification Database (PRIDE) (Perez-Riverol et al., 2019). Raw DIA-MS data were processed with DIA-NN (Demichev et al., 2020) and MaxLFQ (Cox et al., 2014), quantifying 8,498 proteins (***Table S2a, Figure S1c***). The ProCan-DepMapSanger dataset significantly expands the existing molecular characterizations of this broad range of cancer cell line models (***Figure 1b***). Pharmacological screens of anti-cancer drugs tested against this cell line panel were also expanded in this study, representing a 48% increase in the number of unique drugs tested (*n* = 625 drugs and investigational compounds; ***Figure 1c***), with a total of 578,238 half-maximal inhibitory concentration (IC_50_) experimentally measured.

**Figure 1.**
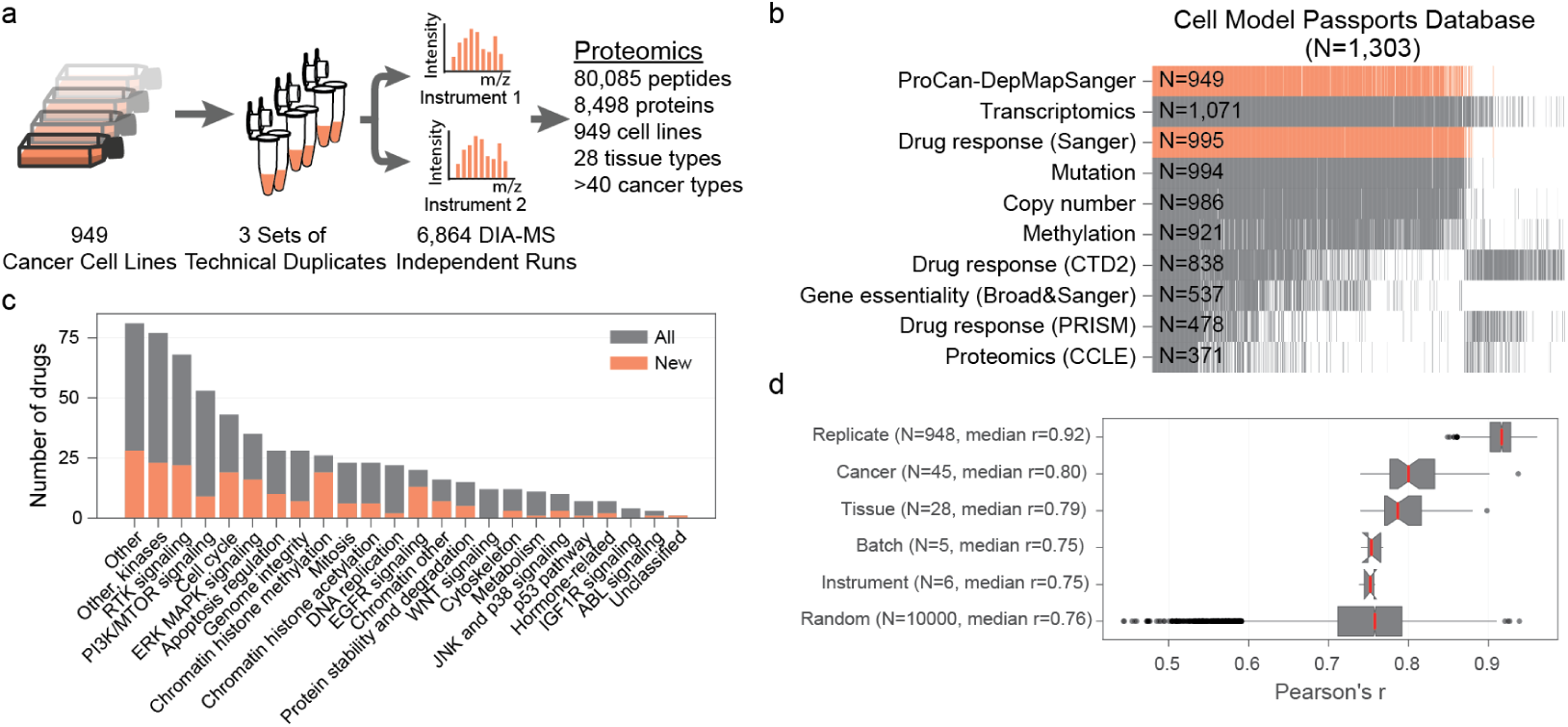
A pan-cancer proteomic map of 949 human cancer cell lines. **a**, Methodology overview for pan-cancer characterization of 949 human cell lines using a DIA-MS workflow. **b**, Proteomic measurements were integrated with independent molecular and phenotypic datasets spanning 1,303 cancer cell lines as part of the Cell Model Passports Database. Data include proteomics (ProCan-DepMapSanger) presented here, transcriptomics, drug response (Sanger), mutation, copy number, methylation, drug response (CTD2), CRISPR-Cas9 gene essentiality (Broad&Sanger), drug response (PRISM), and proteomics (CCLE). Each gray slice denotes a unique cell line, and the total number of cell lines per dataset is indicated. The new proteomic data (ProCan-DepMapSanger) generated in this study are shown in orange, as well as the expanded drug response (Sanger) dataset. **c**, Number of drugs included in the drug response (Sanger) screen, with the orange bar highlighting the additional number of unique drugs presented in this study compared to previous studies. Drugs are grouped by the pathway of their canonical targets. **d**, Pearson’s correlations of the proteomes for each set of six technical replicates, as well as each cancer type, tissue type, batch and instrument. Random indicates the correlation between random unmatched sets of replicates. Median Pearson’s r for each group is reported. Box-and-whisker plots show 1.5 x interquartile ranges and centers indicate medians.

High correlations were observed between replicates of each cell line, yielding a sample-wise median Pearson’s correlation coefficient (Pearson’s *r*) of 0.92 (***Figure 1d*** and ***Figure S1a***). Correlations between unmatched samples from the same instrument or batch were similar to random (median Pearson’s *r* = 0.75, ***Figure 1d***). We also confirmed that our dataset is consistent with other independent previously published proteomic datasets that comprise smaller subsets of the same cell lines (Frejno et al., 2020; Guo et al., 2019; Lawrence et al., 2015; Nusinow et al., 2020; Roumeliotis et al., 2017) (***Figure S1d***). Nonlinear dimensionality reduction using Uniform Manifold Approximation and Projection (UMAP) (McInnes et al., 2018) showed no evidence of instrument or batch effects (***Figure S1e***). Overall, this study generated a high-quality and biologically reproducible pan-cancer proteome map of human cancer cell lines, that is to the best of our knowledge, the largest and broadest of its kind.

### Proteomic profiles reveal cell type of origin

Next we defined a stringent set of protein quantifications that were supported by measuring more than one peptide (*n* = 6,692 human proteins, ***Table S2b***). Visualization of these protein intensities via UMAP showed groupings by cell type of origin (***Figure 2a***). Hematopoietic and lymphoid cells showed the most distinct clustering away from other cell types (***Figure 2a***), and these could be further segregated into different cell lineages (***Figure 2b***). This high-level dimensionality reduction suggested a profile of protein expression that relates to cell type of origin. To investigate this further, a set of 775 proteins that are enriched in certain cell types was selected (***Table S3a***). These cell type-enriched proteins were defined as any protein quantified in > 50% of cell lines from no more than two tissue types, considering only tissues represented by at least 10 cell lines (***Figure 2c***). Cell lines from hematopoietic and lymphoid, peripheral nervous system and skin cell types showed the greatest numbers of these proteins (***Figure 2d***). Using Gene Set Enrichment Analysis (GSEA) (Liberzon et al., 2011; Subramanian et al., 2005), proteins encoded by genes enriched for gene ontology terms for lymphocyte activation, neuron projection and pigmentation were identified in each of these cell types, respectively. Further, the cell type-enriched proteins had a higher correlation between the transcriptome and proteome than did other proteins, suggesting that these represent cell type-specific processes that are more highly conserved between transcription and translation (***Figure 2e***). In support of this, GSEA further identified significant enrichment for genes harboring binding sites for transcription factors related to cell type, including ETS2 in hematopoietic and lymphoid cell lines, KDM7A in peripheral nervous system cell lines and SOX9 in skin cell lines (Fu et al., 2017; Huang et al., 2010; Passeron et al., 2007). Overall, this analysis demonstrates the alignment of the proteomic data with cell lineage, revealing patterns of protein expression that are reflective of cell type of origin.

**Figure 2.**
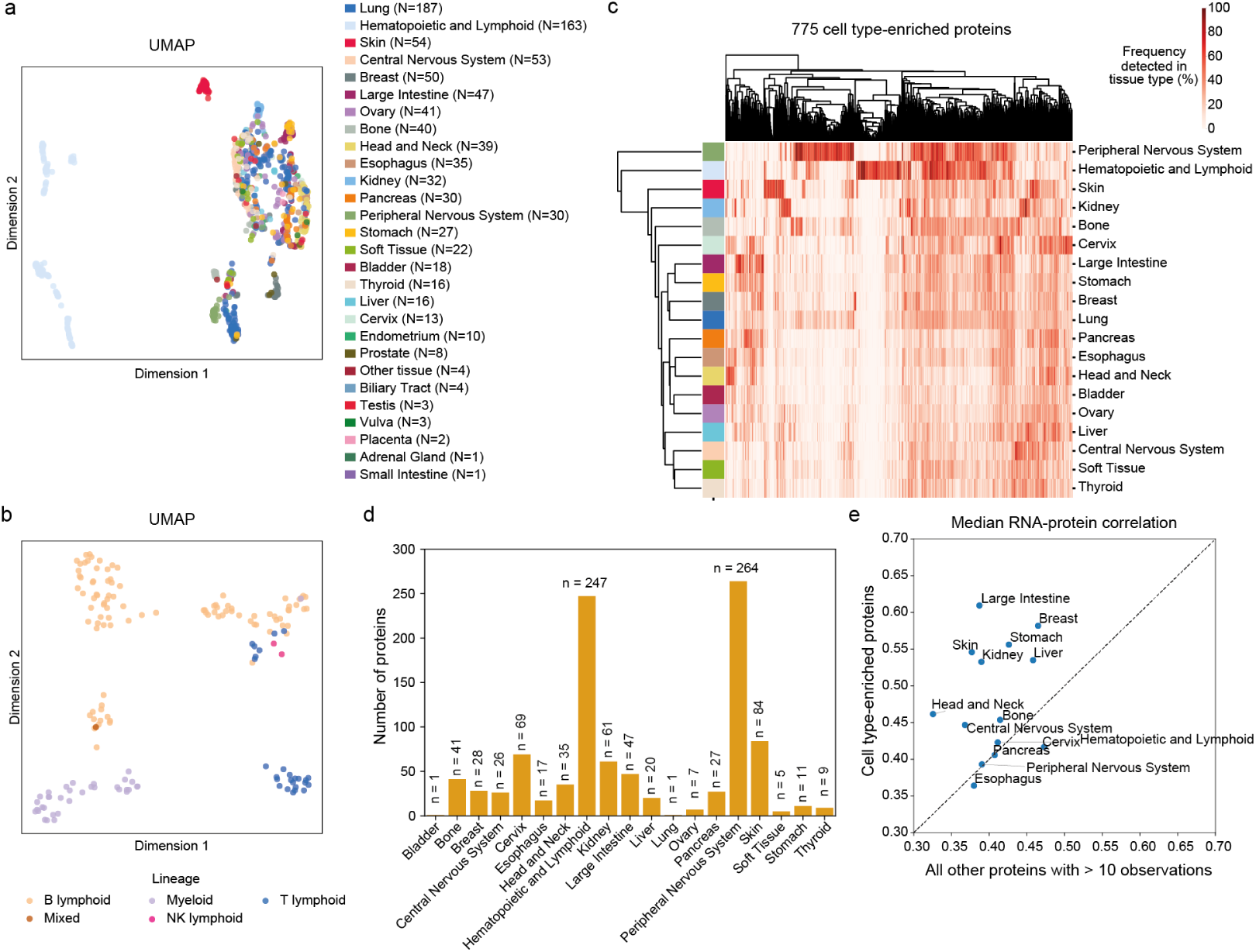
Distinct proteomic profiles according to cell type. **a**, Proteomic data dimensionality reduction by UMAP, with cell lines colored by tissue. **b**, UMAP of hematopoietic and lymphoid cell lines colored by cell lineage. **c**, Heatmap of the frequency of cell type-enriched proteins observed within each tissue. Tissues and proteins are clustered on the vertical and horizontal axes, respectively. **d**, Number of cell type-enriched proteins identified in each tissue type represented by more than 10 cell lines. **e**, Median RNA-protein correlation of cell type-enriched proteins against all other proteins with more than 10 observations in that tissue type. Only tissues with at least 10 cell type-specific proteins are shown.

### Post-transcriptional regulation in diverse cancer cell types

We next sought to identify the key drivers of the distinct protein expression patterns observed across the cell line panel, and to investigate how these integrate with other molecular and phenotypic measurements. Multi-Omics Factor Analysis (MOFA) (Argelaguet et al., 2018, 2020) was used to integrate the proteomic measurements with a range of molecular (promoter methylation, gene expression and protein abundance) and phenotypic (drug response) datasets available for most cancer cell lines (***Table S4*** and ***Figure 3a***). MOFA provides a framework for unsupervised integration of datasets to infer a set of factors (latent variables) that account for biological and technical variability in the data (Argelaguet et al., 2018, 2020). Epithelial-to-mesenchymal transition (EMT) canonical markers, vimentin (VIM) and E-cadherin (CDH1), and EMT GSEA enrichment scores (Liberzon et al., 2011; Subramanian et al., 2005) were found to be associated with the first two factors (F1 and F2) corresponding with large portions of the variability across all datasets (***Figure 3a***). In contrast, technical factors explained little or no variability (***Figure S2a***). Cancer cell lines from the same tissue of origin showed gradients of EMT markers (***Figure 3b***), which are known to be associated with different stages of cancer progression including initiation, metastasis and development of therapy resistance (Brabletz et al., 2018). We observed that some factors capture tissue-specific processes, with their loadings being enriched toward the cell type-enriched proteins defined earlier (***Figure 3a***). For example, MOFA analysis highlighted an association between Factor 12 and skin derived cell lines (***Figure S2b***). Factor 12 also related to phenotypic measurements that are typical of skin cancer cell lines, correlating with CRISPR-Cas9 gene essentiality scores for BRAF (***Figure 3c***) and with its inhibitor, dabrafenib (***Figure 3d***), both of which are strongly associated with cell lines harboring BRAF mutations that are very common in cutaneous melanomas.

**Figure 3.**
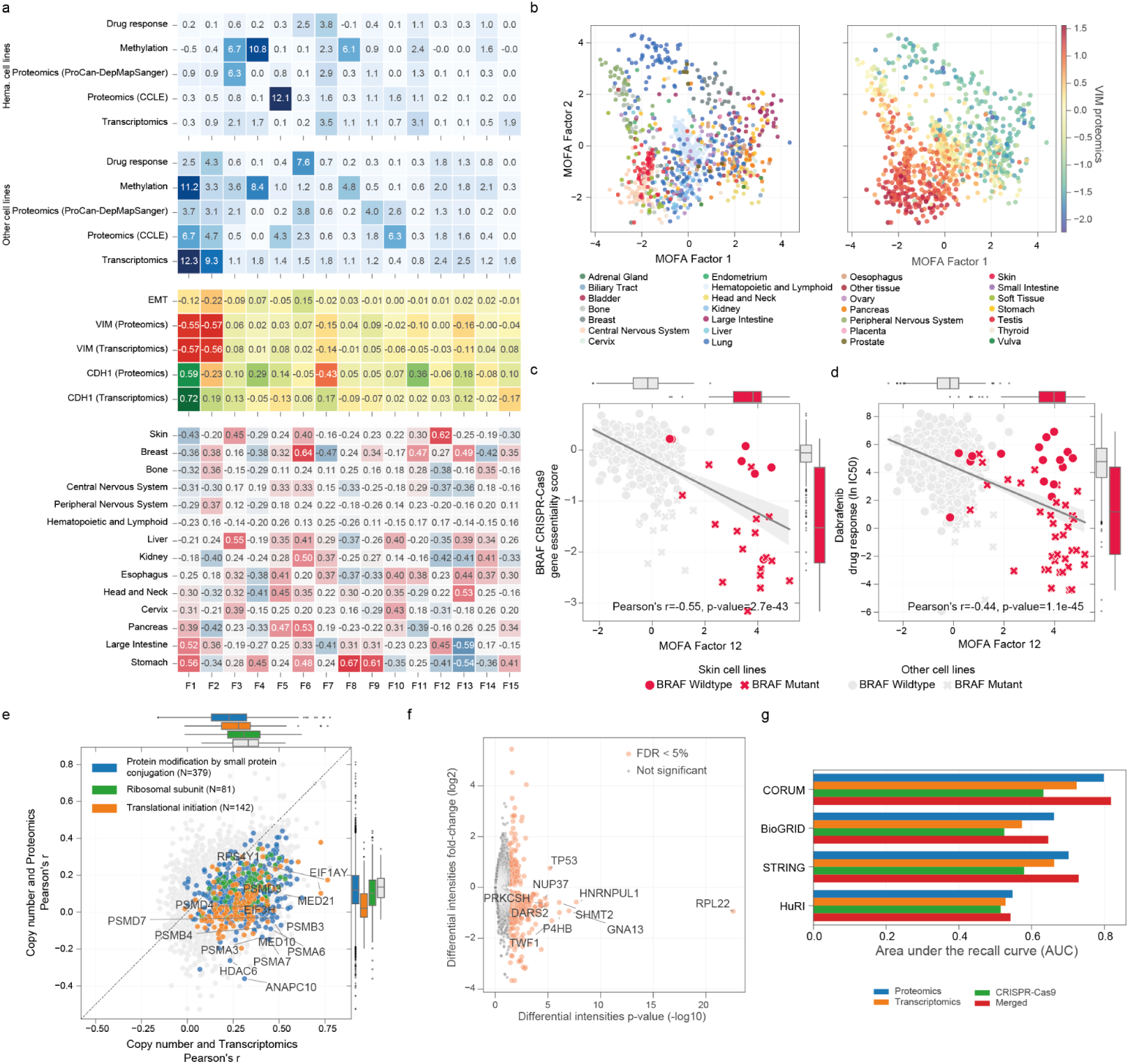
Post-transcriptional regulatory mechanisms of cancer cell lines. **a**, Identification of shared variability (factors) from MOFA across multiple molecular and phenotypic cancer cell line datasets. Hematopoietic and lymphoid cells are grouped and trained separately from the other cell lines. The upper two heatmaps (blue) report the portion of variance explained by each factor (columns) in each dataset. The central (yellow) heatmap reports Pearson’s r between each learned factor and various molecular characteristics of the cancer cell lines. The lower heatmap shows GSEA enrichment scores of each factor to cell type-specific proteins. **b**, Separation of cancer cell lines by MOFA Factors 1 and 2, colored by tissue of origin (left) and by EMT canonical marker VIM protein intensities (right). **c**, Scatter plot with linear regression between MOFA Factor 12 and BRAF CRISPR-Cas9 gene essentiality scores. Skin cancer cell lines are highlighted in red, and BRAF mutant cell lines are marked with a cross. **d**, Similar to **c**, but instead the vertical axis indicates the dabrafenib drug response (IC_50_) measurements. **e**, Pearson’s r between gene absolute copy number profiles with transcriptomics (horizontal axis) and with protein intensities from the ProCan-DepMapSanger dataset (vertical axis). Representative significantly enriched gene ontology terms for proteins with the highest differences between the Pearson’s r are shown, and representative proteins are labeled. N indicates the number of proteins classified by each gene ontology term. Box-and-whisker plots represent the Pearson’s r distributions of proteins involved in each highlighted gene ontology term compared to all proteins (gray). **f**, Volcano plot showing the differential protein intensities between wild-type and mutant cell lines included in the ProCan-DepMapSanger dataset. The top 10 proteins by p-value are annotated. **g**, Recall of protein-protein interactions (PPIs), i.e. ability to detect known PPIs, from resources CORUM, STRING, BioGRID and HuRI. All possible protein pairwise correlations (Pearson’s p-value) were ranked, using proteomics, transcriptomics and CRISPR-Cas9 gene essentiality. The merged score was defined as the product of the p-values of the different correlations.

While EMT was largely concordant between transcriptomics and proteomics, more broadly, a modest and variable association was observed between protein and transcript measurements (median protein-wise Pearson’s *r* = 0.42, ***Figure S2c***). This is consistent with the expectation that proteomic data capture variability explained by post-transcriptional regulation and the proteostasis network. Focussing on the impact of genomic alterations on proteomic profiles, gene copy number was more weakly correlated with protein levels than with gene expression, indicating attenuation of copy number effects between the transcriptome and the proteome (***Figure 3e***). This was particularly evident among subunits of protein complexes such as those involved in ribosomes (***Figure S2d***), which can co-regulate their abundance post-transcriptionally to maintain complex stability and stoichiometry (Gonçalves et al., 2017; Roumeliotis et al., 2017; Ryan et al., 2017; Sousa et al., 2019). Proteins involved in protein synthesis and degradation had some of the strongest post-transcriptional regulation, with several proteasome and ribosome subunits showing strong attenuation (***Figure 3e***). Together, this reflects an active proteostasis network, revealed primarily via direct measurement of the proteome (Gumeni et al., 2017).

Using somatic mutation data to stratify the full set of protein quantifications according to mutation status (***Table S3b***), revealed 478 proteins with significant differential protein intensities between wild type and mutant cell lines (Benjamini-Hochberg adjusted moderated *t*-test < 5%) (***Figure 3f***). When mutations were present in the corresponding gene, the majority of these proteins had decreased abundance (*n* = 354 proteins). In contrast, mutations in *TP53* were associated with significantly higher protein intensities than the wild type (empirical Bayes moderated t-test *p-value* < 0.0001, see ***Methods***). This is consistent with the known increase in stability of many mutant P53 proteins, which results from a decreased rate of proteasome-mediated degradation (Vijayakumaran et al., 2015).

These results indicate that, while variability in protein expression is associated with other molecular and phenotypic layers, there is only partial correlation between transcript and protein abundance, consistent with the effects of post-transcriptional regulation. Thus, the ProCan-DepMapSanger dataset captures additional protein-specific information that can augment our understanding of the impact of genomic alterations affecting, among others, well-established cancer genes.

### Co-regulatory protein networks of cancer cells

Having observed post-transcriptional co-regulation of protein complex abundance (***Figure 3e***), we next investigated whether the abundance of co-regulated proteins could be used to predict putative protein-protein interactions (PPIs). We assessed all possible pairwise protein correlations (*n* = 16,580,952) and as a comparator, used corresponding gene expression and CRISPR-Cas9 gene essentiality profiles, where available. As expected, paralogs and protein complex subunits had some of the strongest correlations (***Table S5***; absolute Pearson’s *r* > 0.5 and false discovery rate (FDR) adjusted *p*-value < 5%). We systematically assessed this enrichment using multiple resources for protein interactions: protein complex interactions from the Comprehensive Resource of Mammalian Protein Complexes (CORUM) (Ruepp et al., 2010); functional interactions from the Search Tool for the Retrieval of Interacting Genes/Proteins (STRING) database (Szklarczyk et al., 2017); physical protein interactions from the Biological General Repository for Interaction Datasets (BioGRID) (Chatr-Aryamontri et al., 2015); and the Human Protein Interactome (HuRI) (Luck et al., 2020). Proteomic measurements had greater ability to detect known PPIs (Area under the recall curve (AUC) = 0.55-0.80) across all resources than transcriptomics (AUC = 0.53-0.72) and CRISPR-Cas9 gene essentiality (AUC = 0.51-0.63) (***Figure 3g***), indicating that PPIs and co-regulation are best captured by proteomics.

The correlation *p*-value score (Fisher’s combined probability test) for BioGRID and HuRI databases was improved slightly by combining different datasets (AUC = 0.54-0.82), suggesting that different types of interactions are captured across multi-omic layers. Proteins with higher numbers of positive protein-protein correlations were more essential for cancer cell survival, as observed in the CRISPR-Cas9 gene essentiality dataset (***Figure S2e***). This is likely linked to their increased transcript and protein expression levels, and the number of pathways in which they are involved. Paralogs were an exception, as they had largely non-essential profiles independent of their gene expression (***Figure S2f***), consistent with their functional redundancy attenuating the impact of loss (Dandage and Landry, 2019).

Considering the high correlations that were observed with known interactions, these pairwise protein correlations could be used to identify novel putative PPIs, such as between protein subunits. Consistent with this, we identified 1,182 putative PPIs with a Pearson’s r greater than 0.8 that are not reported in any of the protein network resources analyzed here. For example, there were strong correlations between protein profiles for EEF2-EIF3I, RPSA-SERBP1 and CCT6A-EEF2 (***Table S5***). These proteins are not reported to interact directly but are also closely related in the high confidence STRING protein interaction network (Szklarczyk et al., 2017). Overall, this analysis highlights the utility of protein measurements for predicting co-regulated interactions.

### Deep learning identifies biomarkers of cancer vulnerabilities

Next, we considered the application of proteomics to identify biomarkers by harnessing drug (Garnett et al., 2012; Gonçalves et al., 2020; Iorio et al., 2016; Picco et al., 2019) and CRISPR-Cas9 gene essentiality (Behan et al., 2019; Meyers et al., 2017; Pacini et al., 2021) screens. The previous Sanger pharmacological screens were expanded to include a total of 578,238 IC_50_ values (***Figure 1b***). These included a total of 625 unique anti-cancer drugs (48% increase in unique drugs) that were screened across 947 of the 949 cancer cell lines, including Food and Drug Administration (FDA)-approved drugs, drugs in clinical development and investigational compounds. To identify protein biomarkers predictive of cancer cell line response to these drugs or CRISPR-Cas9 gene essentialities (*n* = 17,486), we applied linear regression to test all pairwise associations between proteins, drug sensitivity and CRISPR-Cas9 gene dependencies, while considering potentially confounding effects (see ***Methods***) (***Figure 4a***). Among the strongest significant associations (FDR < 10%), we observed many known interactions, including negative associations between ERBB2 protein abundance and gene essentiality scores and MET protein abundance and its inhibitor drug response (***Figure 4b***). Non self-interactions were also observed, such as PPA1-PPA2 paralog synthetic-lethal interaction, where cell lines with lower PPA2 abundance are more sensitive to PPA1 knockout (***Figure S3a***). Moreover, we observed many significant associations between drugs and their known protein targets or functionally related proteins (for example, gefitinib and ERBB2, and Nutlin-3a and BAX) (***Figure 4a***). The majority of the significant drug-protein target associations showed a negative effect size (***Figure 4a***), indicating stronger sensitivity of a cell line to a drug when the target of the drug is more abundant. Lastly, the identification of associations with reported gene copy number alterations, such as amplification of MET and ERBB2, are also observed at the protein level (***Figure 4b***).

**Figure 4.**
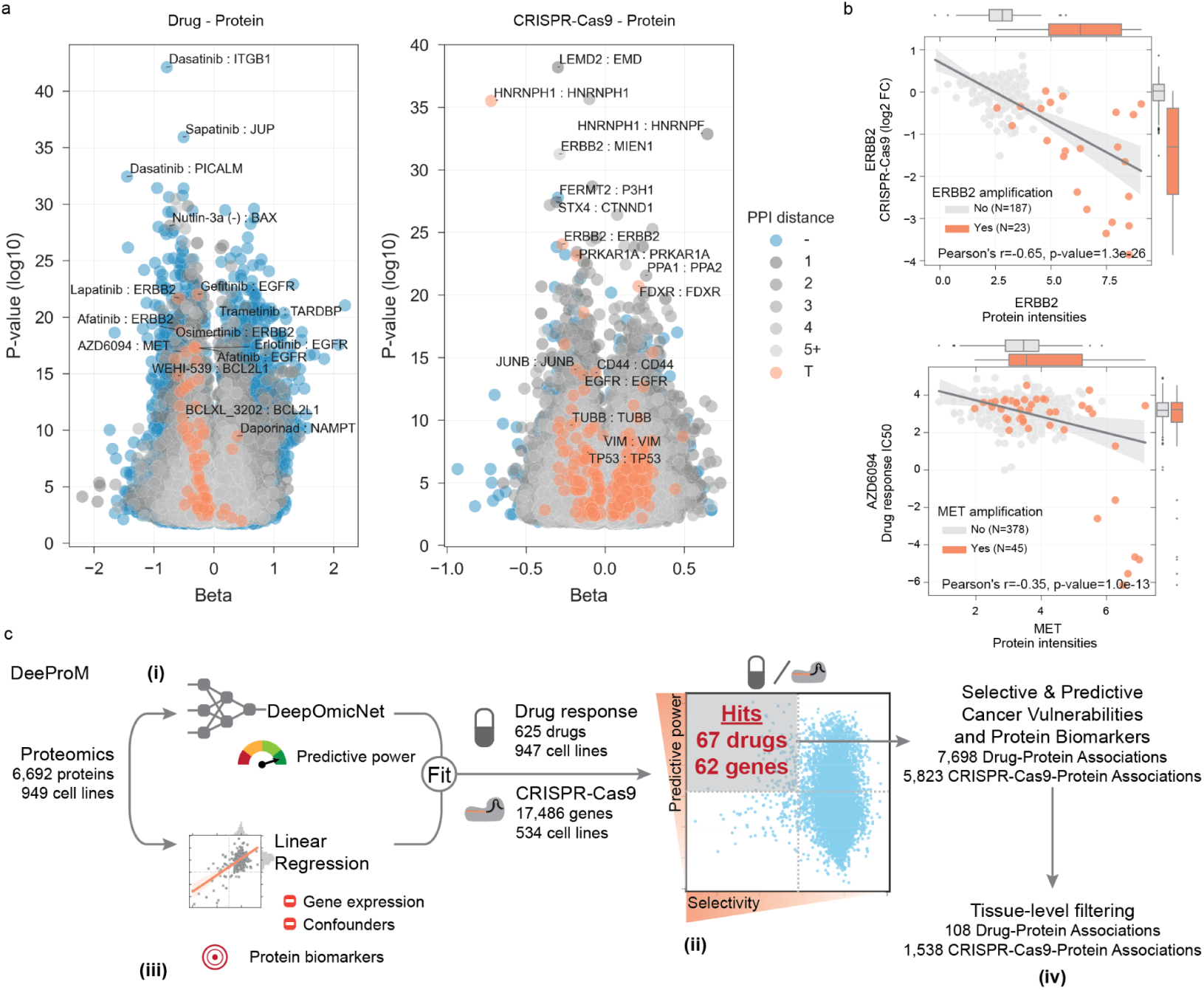
DeeProM identifies protein biomarkers for cancer vulnerabilities. **a**, Significant linear regression associations (FDR < 10%) between protein measurements and drug responses (left panel) and protein measurements and CRISPR-Cas9 gene essentiality scores (right panel). Each association is represented using the linear regression effect size (beta) and its statistical significance (log-ratio test), and colored according to the distance between the target of the drug or CRISPR-Cas9 and the associated protein in a protein-protein interaction network assembled from STRING. T denotes the associated protein is either a canonical target of the drug or the CRISPR-Cas9 reagents; numbers represent the minimal number of interactions separating the drug or CRISPR-Cas9 targets to the associated proteins; and the symbol ‘-’ denotes associations for which no path was found. Representative examples are labeled. **b**, Representative top-ranked CRISPR-Cas9-protein and drug-protein associations. Upper panel shows ERBB2 protein intensities associated with CRISPR-Cas9 gene essentiality, where cell lines with ERBB2 amplifications are highlighted in orange. Lower panel shows the association between AZD6094 MET inhibitor and MET protein intensities, where MET amplified cell lines are highlighted in orange. **c**, Overview of the DeeProM workflow: (i) deep learning models of DeepOmicNet were trained to predict drug responses and CRISPR-Cas9 gene essentialities, prioritizing those that are best predicted by proteomic profiles; and (ii) Fisher-Pearson coefficient of skewness was calculated to identify drug responses and CRISPR-Cas9 gene essentialities that selectively occur in subsets of cancer cell lines. The selected candidates from (i) and (ii) are illustrated by the gray box. (iii) Linear regression models were fitted to identify significant associations between protein biomarkers, drug responses and CRISPR-Cas9 gene essentialities. (iv) Filtering algorithms were applied to further identify tissue-specific cancer vulnerabilities.

To identify biomarker associations that are unique to the proteome and could not be predicted by gene expression measurements alone, we developed a deep learning-based computational pipeline called Deep Proteomic Marker (DeeProM) (***Figure 4c***). DeeProM is powered by DeepOmicNet (see ***Figure S3b*** and ***Methods*** for more details), a deep neural network architecture designed to prioritize drug responses and CRISPR-Cas9 gene essentialities that are highly predictive and specific to subsets of cancer cell lines. As a benchmark, we found that DeepOmicNet consistently outperformed other machine learning approaches, such as elastic net and Random Forest, across a range of multi-omic data sets used in previous studies (Iorio et al., 2016) (***Figure S3c***). In addition, DeeProM incorporates a systematic biomarker search using linear models that take into consideration potential confounding factors, as well as gene expression measurements as covariates (see ***Methods***) to highlight biomarkers that are only evident at the proteomic level. DeeProM assessed all possible drug-protein (*n =* 4,218,788) and CRISPR-protein (*n =* 86,584,537) associations to identify cancer vulnerabilities that are simultaneously well predicted and selective in subsets of cell lines (***Figure 5a***). These two selection criteria (*see* ***Methods***) yielded 67 drug responses and 62 gene essentialities, with a total of 7,698 significant drug-protein and 5,823 significant CRISPR-Cas9-protein associations, that were not significant when considering gene expression measurements alone (***Figure 4c*** and ***Table S6***).

**Figure 5.**
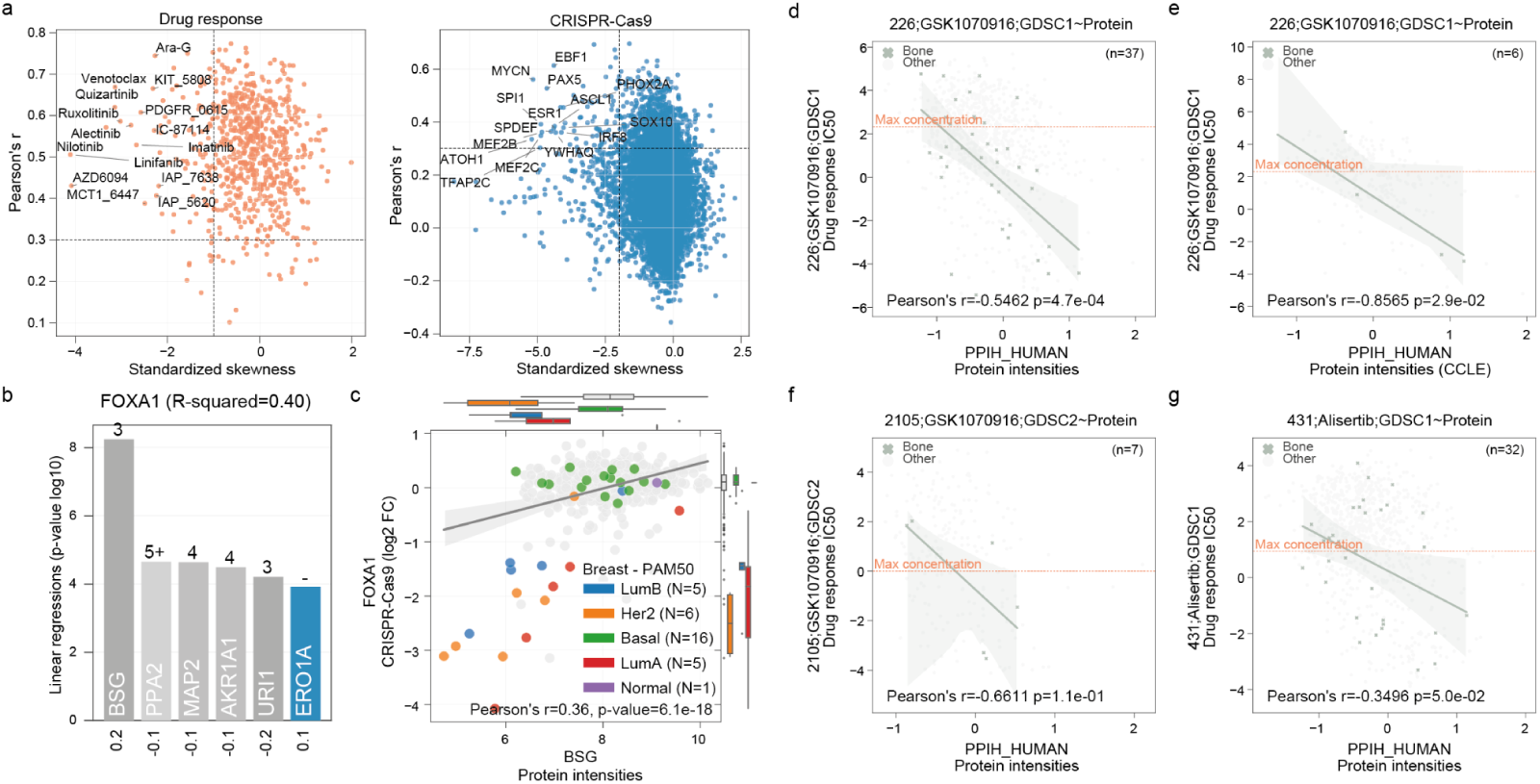
Protein biomarkers identified by DeeProM. **a**, Predictive performance and selectivity of all drug responses (left) and CRISPR-Cas9 gene essentialities (right) across 947 and 534 cancer cell lines, respectively. Data points toward the top left corner of each plot indicate drug responses or gene essentialities that are both selective and well predicted. Top selective drugs and CRISPR-Cas9 gene essentialities are labeled. **b**, Top significant protein associations with FOXA1 CRISPR-Cas9 gene essentiality scores, each bar representing the statistical significance of the linear regression, and below the effect size (beta). The minimal distance of PPIs in the STRING network between FOXA1 and each protein is annotated in each respective bar and color coded according to the description in **Figure 4a. c**, Synthetic-lethal association between FOXA1 CRISPR-Cas9 gene essentiality scores and BSG protein intensities. Breast cancer cell lines are highlighted and sub-classified using the PAM50 gene expression signature (Parker et al., 2009). Box-and-whisker plots indicate the PAM50 subtypes of breast cancers. **d-g**, Representative examples of tissue-specific associations between drug responses and protein markers for cell lines derived from bone tissue (green; all other cell lines are shown in gray). The dashed line represents the maximum concentration used in the drug response screens. **d-f**, The highlighted GSK1070916-PPIH association in bone tissue supported by **d**, the ProCan-DepMapSanger proteomic dataset, **e**, the CCLE proteomic dataset and **f**, the GDSC2 drug response dataset. **g**, Similar to **d**, instead showing data for the drug Alisertib.

Promising targeted therapeutics are often developed for specific cancer types and can display tissue-specific responses. For this reason, DeeProM was used to interrogate associations at the tissue type level by applying a filtering strategy (***Figure 4c*** and *see* ***Methods***). Using the DeeProM workflow, we identified 1,538 tissue type level CRISPR-Cas9-protein associations (***Table S7***). Among the strongest was the dependency on FOXA1 transcription factor knockout and protein levels of basigin (BSG; also known as CD147), a plasma membrane protein expressed in breast cancer cells (***Figure 5b-c***). This association was not observed at the gene expression level (***Figure S3d***). FOXA1–BSG synthetic lethality occurred in luminal (luminal A and B) and HER2-positive (non-basal) breast cancer cell lines, in which BSG protein abundance is low (***Figure 5c***). BSG has been implicated in breast cancer progression (Landras et al., 2019), and is a marker of the aggressive basal-like and triple-negative subtypes, as well as being associated with poor overall survival within these patients (Liu et al., 2018). These data support a model where BSG protein expression is associated with basal-like breast cancer cells, whereas luminal and HER2-positive breast cancer cells, which have low BSG expression, have increased dependency on estrogen receptor-driven FOXA1 transcriptional activity.

DeeProM also identified 108 tissue type level drug-protein associations (***Table S7***). Filtering by the effect size, the strongest association identified was between sensitivity to Aurora kinase B/C inhibitor GSK1070916 and the protein abundance of peptidyl-prolyl cis-trans isomerase H (PPIH) in cell lines derived from bone tissue (***Figure 5d***). This association was significant at the protein level but was not significant in the transcriptome (***Figure S4a***). The association was further supported by examination of the Cancer Cell Line Encyclopedia (CCLE) proteomic dataset (***Figure 5e*** *and* ***Figure S4b***) (Nusinow et al., 2020), using an independent screening of GSK1070916 in the Sanger drug sensitivity dataset (***Figure 5f*** *and* ***Figure S4c***), and with the PRISM drug response dataset (***Figure S4d***) (Corsello et al., 2020), in which there was a suggestive association that did not reach statistical significance due to the smaller sample size. Furthermore, there was a strong association between PPIH protein levels and Alisertib, a second Aurora kinase inhibitor (***Figure 5g*** and ***Figure S4e-f***). PPIH and Aurora kinase A are both regulated by the p53-p21-DREAM-CDE/CHR signaling pathway (Fischer et al., 2016), supporting the identified link between Aurora kinase inhibitor sensitivity and PPIH protein levels.

Taken together, these results demonstrate the unique value of proteomic measurements for the discovery of cancer biomarkers. We identified both established and novel cancer related biomarkers, including protein biomarkers for selective cancer vulnerabilities that cannot be found using gene expression measurements alone.

### Predictive power of protein sub-networks on cancer cell phenotypes

We have established the utility of proteomics to identify specific biomarkers for cancer vulnerabilities. Using cell lines also included in an independent proteomic dataset from the CCLE (Nusinow et al., 2020), we observed comparable performance to the ProCan-DepMapSanger dataset when predicting three independent drug response datasets (***Figure 6a*** and ***Figure S5a***) and CRISPR-Cas9 gene essentiality profiles (***Figure 6b*** and ***Figure S5b***). We next compared the predictive power of proteomic and transcriptomic data for modeling drug responses and CRISPR-Cas9 gene essentialities. The predictive power of our models was highly similar when trained using either the ProCan-DepMapSanger or transcriptomics dataset (***Figure 6c***). This was recapitulated by machine learning methods such as elastic net and Random Forest (***Figure 6d***). Notably, the predictive performance of protein measurements alone outperformed the corresponding overlapping subset of the transcriptome (***Figure S5c***). Proteomic measurements further showed overall stronger protein pairwise correlations than transcriptomics or CRISPR-Cas9 gene essentialities (***Figure 6e***). This demonstrates that proteomics can provide additional relevant information that is not captured by transcriptomics.

**Figure 6.**
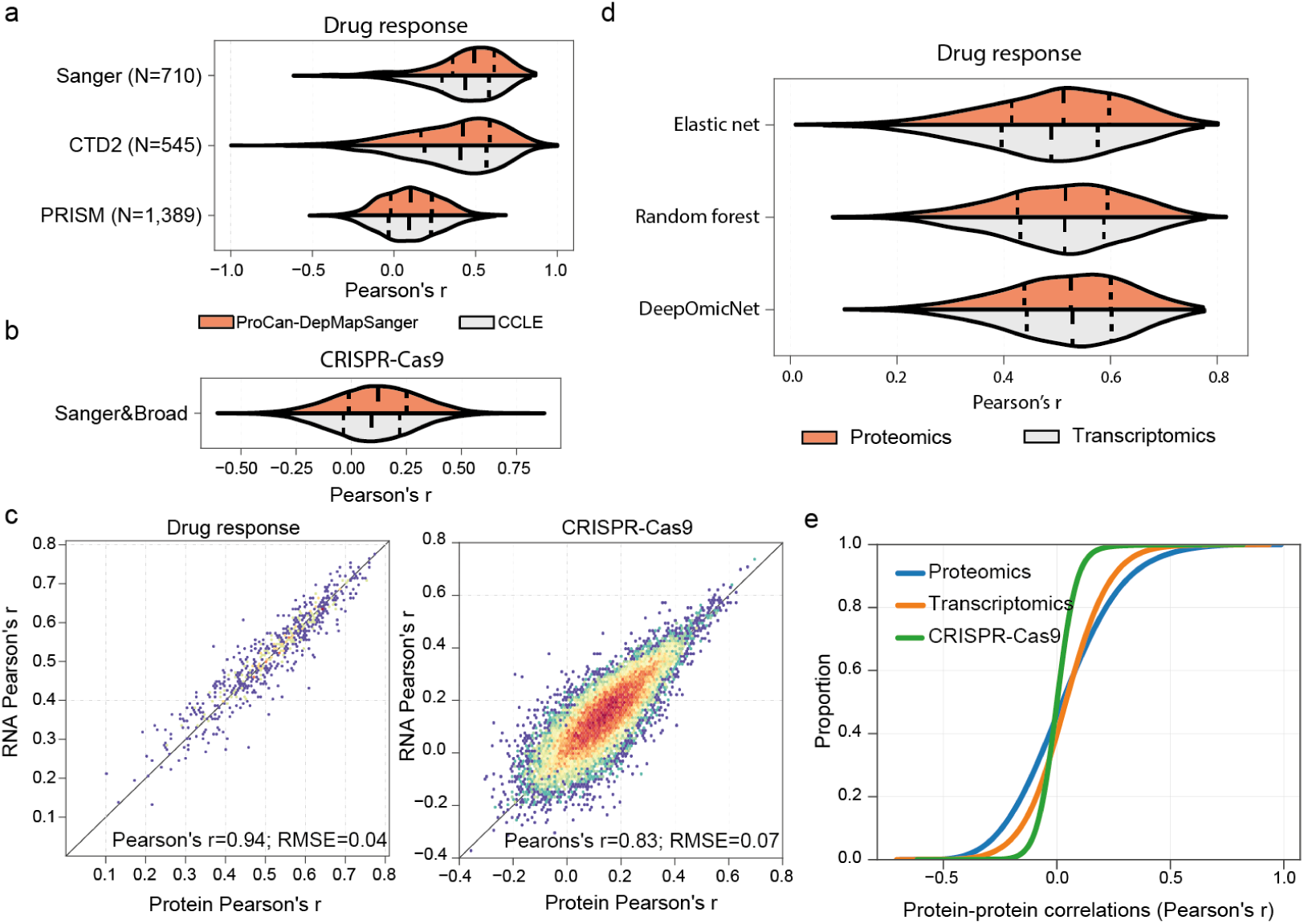
Evaluation of predictive power of DeepOmicNet for multi-omic datasets. **a-b**, Distribution of the predictive power (mean Pearson’s r between predicted and observed IC_50_ values) of DeepOmicNet, comparing ProCan-DepMapSanger to an independent proteomic dataset (CCLE), using cell lines in common between the two datasets. Plots show prediction of **a**, drug responses (N represents the total number of drugs tested; n = 290 cell lines) and **b**, CRISPR-Cas9 gene essentialities (n = 234 cell lines). **c**, Two-dimensional density plots showing the predictive power of DeepOmicNet in predicting drug responses (left) and CRISPR-Cas9 gene essentiality profiles (right) using protein (horizontal axis) and transcript (vertical axis) measurements. Each data point denotes the Pearson’s r between predicted and observed measurements for each drug or CRISPR-Cas9 gene essentialities. **d**, Similar to **a**, distribution of the predictive power of three machine learning models using either proteomic or transcriptomic measurements to train and predict drug responses (Sanger dataset). **e**, Cumulative distribution function of the Pearson’s r of all pairwise protein-protein correlations compared with transcriptomics and CRISPR-Cas9 gene essentiality measurements.

To determine how predictive power is influenced by the number of proteins quantified, a random downsampling analysis was performed to predict drug responses, with a decrease of 500 proteins in each step (see ***Methods***). This showed that a randomly selected subset of 1,500 proteins was able to provide 88% of the predictive power of the full dataset (mean Pearson’s *r* = 0.43 at *n* = 1,500 proteins versus mean Pearson’s *r* = 0.49 at *n* = 8,498 proteins; ***Figure 7a***). This implies that a fraction of the quantifiable proteins is sufficient to represent fundamental elements of the proteome involved in mediating key cellular phenotypes, presumably because proteins are organized into complexes with connected and co-regulated subunits. (***Figure 7b***).

**Figure 7.**
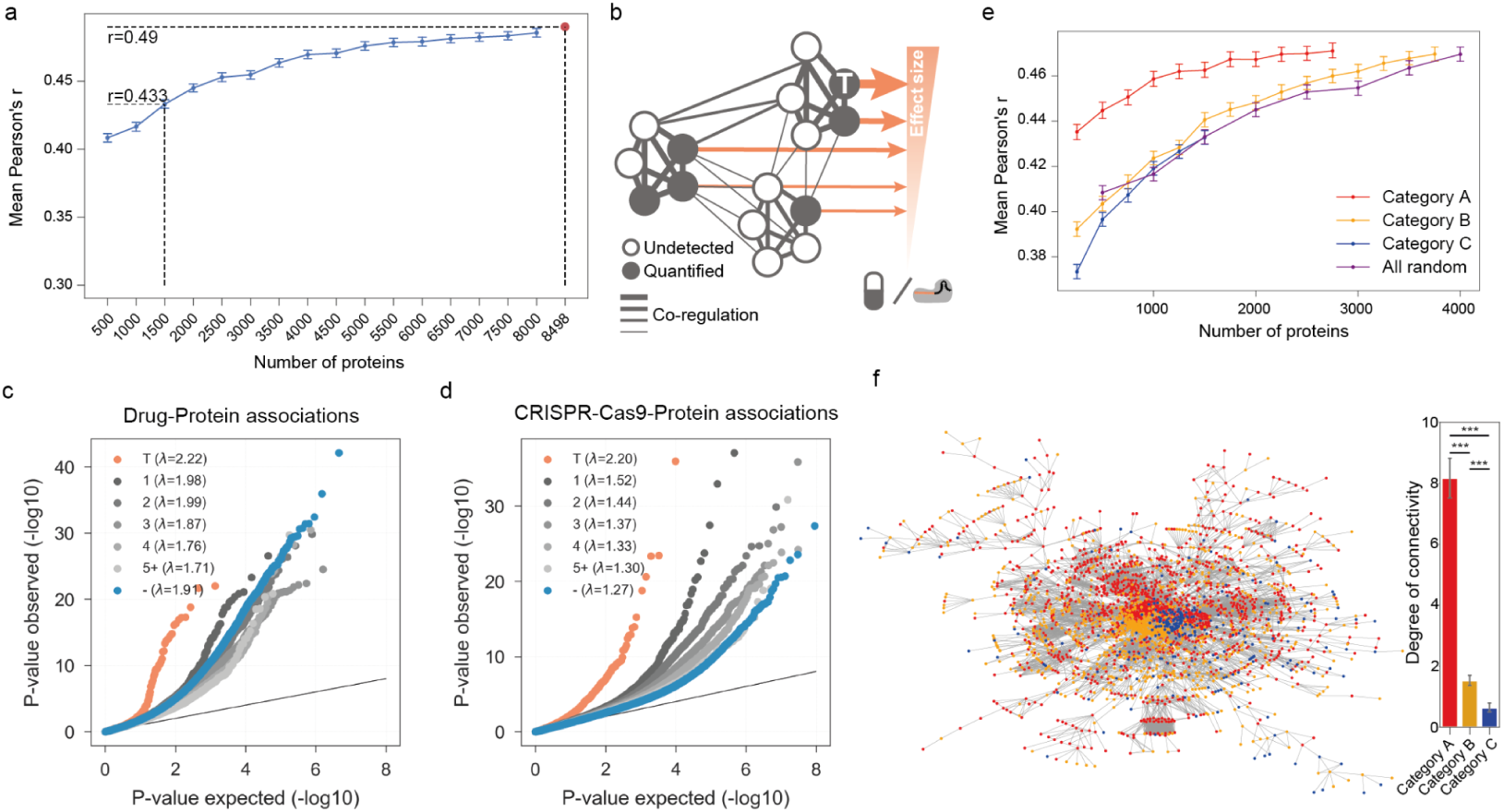
Proteomic support for a network pleiotropy model. **a**, Comparison of the predictive power of DeepOmicNet trained with randomly downsampled sets of proteins. The dots indicate the means and vertical lines represent 95% confidence intervals derived from 10 iterations of random downsampling. The red point represents the full predictive power using all of 8,498 quantified proteins. **b**, Schematic diagram depicting protein network pleiotropy with widespread protein associations with responses to either drugs or CRISPR-Cas9, and demonstrating the strongly co-regulated nature of protein networks. Nodes represent proteins that could be either quantified or are undetected, where ‘T’ represents a protein target of a drug or CRISPR-Cas9 gene essentialities. Edges showcase putative interactions, with high correlation coefficients between proteins depicted by thicker edges. Orange arrows represent the variability explained by that protein for the cancer cell line’s response to a drug or CRISPR-Cas9 gene perturbation. The size of the arrow is proportional to the variance explained. **c-d**, Quantile-quantile plots of protein associations with **c**, drug responses and **d**, CRISPR-Cas9 gene essentiality profiles. Protein associations are grouped and colored by their distances from the drugs or CRISPR-Cas9 targets using the STRING protein interaction network. P-values were calculated in likelihood ratio tests on all parameters of the linear regression models. Annotation is as described in **Figure 4a**. P-value lambda ‘λ’ inflation factors are reported for each drug and CRISPR-Cas9 gene association. **e**, Comparison of the predictive power of DeepOmicNet models trained with subsets of Category A, B and C proteins (per **Figure S5f**) comprising randomly downsampled sets of proteins. The dots indicate the means and vertical lines represent 95% confidence intervals derived from 10 iterations of random samplings. **f**, The STRING protein interaction network diagram, with proteins colored according to Category (left). The bar chart (right) shows the network connectivity for these proteins, where degree represents the number of other proteins connected to a given protein according to the STRING PPI network. *** denotes significant at P < 0.001 by unpaired t-test.

To investigate protein networks, DeeProM analyses of drug-protein and CRISPR-Cas9-protein associations were examined in the context of protein networks (***Figure 7c-d***). The strongest overall associations were observed between the drug and CRISPR-Cas9 targets and their protein intensities (***Figure 7c-d***). Additionally, CRISPR-Cas9-protein associations showed that proteins closer in the PPI network to the targeted proteins had stronger associations than those further apart (***Figure 7d***). This enrichment for targets and functionally closer proteins remains even when the contribution of transcriptomic measurements is removed for the drug-protein and CRISPR-Cas9-protein associations (***Figure S5d-e***). However, many seemingly functionally distant proteins (> 2 steps away from the perturbation target in PPIs) also exhibit significant drug-protein and CRISPR-Cas9-protein associations (***Figure 7c-d***).

We next explored the relationship between predictive performance and sub-networks comprising proteins of differing frequencies in the full dataset, defining categories of proteins as those found in >= 90% (Category A; *n* = 2,944 proteins), 20-90% (Category B; *n* = 3,939) or < 20% (Category C; *n* = 1,615) of the cell lines, respectively (***Figure S5f***). Downsampling these protein sets at random, with a decrease of 250 proteins in each step (see ***Methods***), showed that proteins that are frequently observed in the dataset (Category A), provided the highest predictive performance (for example, mean Pearson *r* = 0.463 for 1,500 proteins, or *r* = 0.435 for 250 proteins) when compared against less frequently observed proteins (Category B, mean *r* = 0.441; and Category C, mean *r* = 0.432, for 1500 proteins; ***Figure 7e***). Category A proteins had a significantly higher degree of connectivity (mean of 8 degrees) than Category B and C proteins (mean of 2 and 1 degree(s), respectively) in the STRING protein interaction network (Szklarczyk et al., 2017) (***Figure 7f***). Together, these results demonstrate that quantification of small subsets of commonly expressed proteins within highly-interconnected networks can be used for predictive modeling of cellular phenotypes.

## Discussion

The ProCan-DepMapSanger data resource is the largest pan-cancer proteomic map available to date and provides multiple new insights beyond existing molecular datasets. This map quantifies 8,498 proteins across 949 human cancer cell lines, representing 28 tissues and over 40 histologically diverse cancer types and a wide range of genotypes, significantly expanding the molecular characterization of cancer models as part of a Cancer Dependency Map (Boehm et al., 2021). All data are publicly available along with other molecular and phenotypic datasets at http://cellmodelpassports.sanger.ac.uk (van der Meer et al., 2019). This proteomic dataset is a high-quality resource for mechanistic investigation of network organization and regulatory principles of the proteome, as well as for translational discoveries.

This study demonstrated protein expression patterns that reflect the epithelial-to-mesenchymal transition, and molecular distinctions according to cell lineage. The data also revealed widespread protein regulatory events, such as post-transcriptional attenuation of gene copy number effects. ProCan-DepMapSanger allowed the comprehensive characterisation of protein expression patterns that could not be captured by the transcriptome, exposing the benefits of directly measuring protein abundance. Furthermore, we developed a novel deep learning-based pipeline DeeProM, with a deep neural network architecture, which consistently outperformed other machine learning approaches. DeeProM enabled the full integration of proteomic data with drug responses and CRISPR-Cas9 gene essentiality screens to build a comprehensive map of protein-specific biomarkers of cancer vulnerabilities that are essential for cancer cell survival and growth. Notably, to the best of our knowledge, this is the first comprehensive demonstration that proteomic data spanning a broad range of cancer cell types and molecular backgrounds has significant utility for predicting cancer cell vulnerabilities.

Our proteomic workflow was devised for clinical applicability by using shortened preparation times, low peptide loads and short LC/MS run-time. This has enabled analysis of large numbers of very small cancer samples, with high throughput and minimal instrument downtime. The CCLE proteomic dataset was generated using higher peptide loads and longer LC/MS run-time to measure more proteins (12,755 proteins), and consequently is less compatible with clinical workflows. Despite the different depths of protein coverage, the CCLE and ProCan-DepMapSanger proteomic datasets had equivalent power for predicting cancer dependencies. Similarly, the ProCan-DepMapSanger proteomic dataset had similar predictive power to cancer cell line transcriptomic data. Taken together, this demonstrates that a high-throughput and clinically-relevant sample workflow, as used in this study, produces data with sufficient predictive power to determine cancer dependencies, and indicates the potential of proteomics for clinical applications using small biopsies of human cancer tissue in diverse molecular contexts. Subsequent application of the clinical proteomics sample workflow and integration of the ProCan-DepMapSanger dataset with cancer tissue samples is likely to provide numerous potential clinical applications, such as the proteomic molecular identification and stratification of cancer subtypes.

Measuring even a fraction of the proteome, as small as 1,500 randomly selected proteins, provided power to predict drug responses that was similar to the full proteome that we report. This suggests that random subsets of protein data comprising a relatively small number of proteins would be sufficient to represent many fundamental cellular processes. This is consistent with an omnigenic model (Boyle et al., 2017), whereby large numbers of genes are related to many different disease traits in an interconnected manner. This is related to the proteostasis network model of sustaining proteome balance via coordinated protein synthesis, folding, conformation and degradation (Gumeni et al., 2017). In the context of the cancer proteome, we propose that pleiotropic networks of highly-connected and co-regulated proteins contribute toward establishing cellular phenotypes. This includes a small number of core protein modules that are proximal to the phenotype and have the strongest effect, and a much larger set of more distal peripheral proteins that together explain a significant portion of total variation.

In conclusion, this dataset represents a major resource for the scientific community, for biomarker discovery and for the study of fundamental aspects of protein regulation that are not evident from existing molecular datasets. This will enable the identification of targets and treatments for application in precision oncology approaches.

## Acknowledgments

We thank Ricard Argelaguet for helpful comments and discussions on the implementations of the MOFA analysis, and Keith Ashman for discussions on optimizing the bank of mass spectrometers. ProCan^®^ is supported by the Australian Cancer Research Foundation, Cancer Institute New South Wales (NSW) (2017/TPG001, REG171150), NSW Ministry of Health (CMP-01), The University of Sydney, Cancer Council NSW (IG 18-01), Ian Potter Foundation, the Medical Research Futures Fund (MRFF-PD), National Health and Medical Research Council (NHMRC) of Australia European Union grant (GNT1170739, a companion grant to support the European Commission’s Horizon 2020 Program, H2020-SC1-DTH-2018-1, ‘iPC - individualizedPaediatricCure’ [ref. 826121]), and National Breast Cancer Foundation (IIRS-18-164). The work at ProCan^®^ was done under the auspices of a Memorandum of Understanding between Children’s Medical Research Institute and the U.S. National Cancer Institute’s International Cancer Proteogenome Consortium (ICPC). R.C.P. and P.J.R. are supported by NHMRC Fellowships (GNT1138536 and GNT1137064, respectively). E.G.’s work is supported by UIDB/50021/2020 (INESC-ID multi-annual funding). This research was funded in whole, or in part, by the Wellcome Trust Grant 206194. For the purpose of Open Access, the authors have applied a CC BY public copyright license to any Author Accepted Manuscript version arising from this submission.

## Author contributions

Conceptualization: PJR, QZ, MJG, RRR

Methodology: EG, RCP, ZC, PGH, PJR, QZ, MJG, RRR

Software: EG, RCP, ZC

Formal analysis: EG, RCP, ZC, QZ

Investigation: EG, RCP, ZC, SSM, NL, AB, DBN, CH, JK, SM, IM, JM, LR, AJS, ES, FT, SGW, YW, DX, KLM, QZ

Data curation: SB, MD, MH, KLM, BT

Project administration: SV

Writing – original draft: EG, RCP, ZC, PJR, QZ, MJG, RRR

Writing – review & editing: All authors.

## STAR Methods

### Cancer cell line authentication

The 949 human cancer cell lines used in this study have been obtained from public repositories and private collections and are maintained according to the suppliers’ guidelines. More detailed information about each cell line can be found at the Cell Model Passports portal (https://cellmodelpassports.sanger.ac.uk/) (van der Meer et al., 2019). All cell line stocks were tested for mycoplasma contamination prior to banking using both a polymerase chain reaction (EZ-PCR Mycoplasma Detection Kit, Biological Industries) and a biochemical test (MycoAlert, Lonza). Cultures testing positive using either method were removed from the collection.

To prevent cross-contamination or misidentification, all banked cryovials of cell lines were analyzed using a panel of 94 single nucleotide polymorphisms (SNPs) (Garnett et al., 2012) (Fluidigm, 96.96 Dynamic Array IFC). The data obtained were compared against a set of reference SNP profiles that have been authenticated by short tandem repeat (STR) back to a published reference (typically the supplying repository). Where a published reference STR profile is not available, the reference SNP profile is required to be unique within the collection/dataset. A minimum of 75% of SNPs is required to match the reference profile for a sample to be positively authenticated.

Additionally, one of the replicate cell pellets generated from each cell line for this study underwent authentication via STR profiling at CellBank Australia (Westmead, Australia). To do so, STR loci were amplified using the PowerPlex® 16HS System (Promega) and the data were analyzed using GeneMapper™ ID software (ThermoFisher). Only cell lines that passed this quality control metric were retained for analysis (*n* = 949).

### Cell culture and harvesting

For each cell line, distinct cell pellets from a single cell culture were produced as technical replicates. Cells were cultured to semi-confluence at 37 °C and 5% CO_2_ in the appropriate medium and then harvested. Suspension cells were centrifuged at 200 g for 5 min at 4 °C and the supernatant was removed. The cells were then washed twice by resuspension in ice-cold Dulbecco’s phosphate buffered saline containing no calcium or magnesium (DPBS) and centrifugation at 200 g for 5 min at 4 °C. For adherent cells, the culture medium was removed before washing with ice-cold DPBS and the cells were removed by mechanical scraping into fresh ice cold DPBS. The harvested cells were then centrifuged as before, washed twice in ice cold DPBS, transferred to 1.5 mL centrifuge tubes (Protein LoBind Tubes, Eppendorf) and centrifuged at 600 g for 5 min at 4 °C. The DPBS was removed and the tubes containing the cell pellets were snap frozen on dry ice, then stored at -80 °C.

### Acquisition of mass spectrometry (MS) data

#### Cell lysis and digestion

Three cell pellets were analyzed for each of the 949 cell lines. The cell pellets were processed using Accelerated Barocycler Lysis and Extraction (ABLE) protocol with minor modifications (Lucas et al., 2019). In brief, all cell pellets were centrifuged to remove residual DPBS, then resuspended in a volume of 1% (w/v) sodium deoxycholate (SDC) that was appropriate for the cell count (between 50–400 μL). To this, 1 unit of benzonase was added to digest the DNA/RNA in the samples for 5 min at 37 °C and mixed with shaking at 1000 rpm. After incubation, a 50 µL aliquot was taken and further processed, with peptide digestion carried out as previously published (Lucas et al., 2019).

#### Data Independent Acquisition (DIA)-MS Acquisition

We used a clinically-relevant workflow that enables high throughput and minimal instrument downtime; 2 µg of peptide was loaded for each replicate with 90-minute acquisitions. Three technical replicates of peptide preparations were generated. Each replicate was injected on two of six different SCIEX™ 6600 TripleTOF^®^ mass spectrometers coupled to Eksigent nanoLC 425 high-performance liquid chromatography (HPLC) systems, housed at a single facility, ProCan^®^ in Westmead, Australia (***Figure S1b***). In each case, an Eksigent nanoLC 425 HPLC system (Sciex) operating in microflow mode was coupled online to a 6600 TripleTOF^®^ system (Sciex) run in sequential windowed acquisition of all theoretical fragment ion spectra (SWATH™) mode using 100 variable isolation windows (***Table S8a***). The parameters were set as follows: lower m/z limit 350; upper m/z limit 1250; window overlap (Da) 1.0; collision energy spread was set at 5 for the smaller windows, then 8 for larger windows; and 10 for the largest windows. MS/MS spectra were collected in the range of m/z 100 to 2000 for 30 ms in high resolution mode and the resulting total cycle time was 3.2 s.

The peptide digests (2 µg) were spiked with retention time standards and injected onto a C18 trap column (SGE TRAPCOL C18 G203 300 µm x 100 mm) and desalted for 5 min at 10 µL/min with solvent A (0.1% [v/v] formic acid). The trap column was switched in-line with a reversed-phase capillary column (SGE C18 G203 250 mm × 300 µm ID 3 µm 200 Å), maintained at a temperature of 40 °C. The flow rate was 5 µL/min. The gradient started at 2% solvent B (99.9% [v/v] acetonitrile, 0.1% [v/v] formic acid) and increased to 10% over 5 min. This was followed by an increase of solvent B to 25% over 60 min, then a further increase to 40% for 5 min. The column was washed with a 4 min linear gradient to 95% solvent B held for 5 min, followed by a 9 min column equilibration step with 98% solvent A. The TripleTOF^®^ 6600 system was equipped with a DuoSpray source and 50 µm internal diameter electrode and controlled by Analyst 1.7.1 software. The following parameters were used: 5500 V ion spray voltage; 25 nitrogen curtain gas; 100 °C TEM, 20 source gas 1, 20 source gas 2.

### Spectral library generation and DIA-MS Data processing

An *in silico* spectral library was created using DIA-NN (version 1.8) (Demichev et al., 2020) for the canonical human proteome (Uniprot Release 2021_03; 20,612 sequences), along with retention time peptides and commonly occurring microbial and viral sequences. DIA-MS data in wiff file format were collected for 6,981 MS runs (***Table S8b***), and all of these MS runs were used to create a spectral library in DIA-NN (Demichev et al., 2020). To reduce the search space, the library was confined to precursors identified in the *in silico* library only. The final spectral library contained a total of 12,487 proteins and 144,578 precursors. DIA-NN (version 1.8) (Demichev et al., 2020) was used to process the MS data using this spectral library, implemented using RT-dependent normalization with parameters given in ***Table S8c***. All MS runs, as well as the FASTA and spectral library files, have been deposited in the Proteomics Identification Database (PRIDE) (Perez-Riverol et al., 2019).

DIA-NN output data were filtered to retain only precursors from proteotypic peptides with Global.Q.Value ≤ 0.01. Proteins were then quantified using maxLFQ with default parameters (Cox et al., 2014) and implemented using the DiaNN R Package (https://github.com/vdemichev/diann-rpackage). Data were then log_2_-transformed. 117 files were discarded from downstream analyses (***Table S8b***), as follows: one MS run recorded no peptides, six replicates of SW900 were removed because the cell line failed STR profiling; 32 files from the earliest pilot batch were removed, as these were repeated later in the experiment; 39 files that quantified fewer than 2,000 proteins were removed; 39 files were removed because they had a poor correlation across replicates. Cell lines with a poor replicate correlation were identified using two methods. First, the minimum correlation between replicates was calculated for each cell line. The 10% of cell lines with the lowest correlation across the cohort were then examined to identify whether any MS run had a correlation with an MS run from another cell line that was above the 75% percentile of correlations (*n* = 11 cell lines). MS runs were then discarded for each cell line if manual examination of replicate correlations indicated that a sample mix up could have occurred (*n* = 21 MS runs discarded). Second, any cell line was selected that had a minimum correlation in at least one replicate of < 0.8 or a coefficient of variation, from proteins observed in > 80% of the cohort, across replicates of > 30% (*n* = 9 cell lines). These MS runs were then also manually examined for each cell line and MS runs that were discordant with the remainder of replicates were removed (*n* = 18 MS runs discarded). The final dataset, termed ProCan-DepMapSanger, was derived from 6,864 mass spectrometry runs (***Table S1b***) covering 949 cell lines (***Table S1a***) and quantifying a total of 8,498 proteins (***Table S2a***). A filtering was applied to identify protein quantifications derived from more than one supporting peptide (*n* = 6,692 human proteins; ***Table S2b***). MS runs across replicates of each cell line were combined by calculating the geometric mean. Protein quantifications and number of peptides identified per protein in each MS run are available in PRIDE (Perez-Riverol et al., 2019).

### Assembly of multi-omics cancer cell line datasets

Drug response measurements were assembled from multiple studies (Garnett et al., 2012; Gonçalves et al., 2020; Iorio et al., 2016; Picco et al., 2019) and 204 new compounds were screened and dose response curves fitted as previously described in detail (Iorio et al., 2016; Vis et al., 2016). A total of 625 unique drugs were included in our drug response dataset. All data and respective details can be accessed at www.cancerRxgene.org (Yang et al., 2013). Cell line growth rates were represented as the ratio between the mean of the untreated negative controls measured at day one (time of drug treatment) and the mean of the dimethyl sulfoxide (DMSO) treated negative controls at day four (72 h post drug treatment). Data acquisition and processing was performed as previously described (https://www.cancerrxgene.org/) to systematically fit drug response curves and derive half-maximal inhibitory concentration (IC_50_) measurements for each drug across the cell lines measured (Garnett et al., 2012; Gonçalves et al., 2020; Iorio et al., 2016; Picco et al., 2019; Yang et al., 2013). The dataset comprises two different screening approaches (Yang et al., 2013), and for drugs screened with both modalities, these were kept as separate entries for the downstream analyses by constructing a unique identifier (drug_id) with the pattern of <drug_code>_<drug_name>_<GDSC_version>, resulting in 819 drug_ids. A threshold of a minimum of 300 cell lines was applied to exclude drugs that were screened without enough cell lines for DeeProM analysis, resulting in a total of 710 drug_ids. The natural log of the raw IC_50_ was used for all computations.

RNA sequencing (RNA-seq) transcriptomics and Infinium HumanMethylation450 methylation measurements for the same set of cancer cell lines were assembled from previous analyses, for which the acquisition and processing are described in detail (Garcia-Alonso et al., 2018; Iorio et al., 2016). Mutation and copy-number calls were inferred from whole-exome sequencing and Affymetrix SNP6 arrays, respectively, as described previously (Iorio et al., 2016).

Genome-wide essentiality measurements for 17,486 genes were assembled for 534 cancer cell lines that overlap with those analyzed in the ProCan-DepMapSanger dataset, using CRISPR-Cas9 screens (Pacini et al., 2021). This is an integrated CRISPR-Cas9 dataset derived from two projects (Behan et al., 2019; Meyers et al., 2017) that removes library biases and represents gene essentiality as log_2_ fold-changes corrected for copy number bias (Iorio et al., 2018).

All of these datasets were integrated and made available to query and download through Cell Model Passports data portal (https://cellmodelpassports.sanger.ac.uk/) (van der Meer et al., 2019).

### Dimensionality reduction and visualization

Uniform Manifold Approximation and Projection (UMAP) (McInnes et al., 2018) was calculated using Python package umap-learn (v.0.4.2) with the default setting of 15 nearest neighbors and the first 50 principal components derived from the protein matrix. Missing values were replaced with zero to calculate the principal components with the Python package scikit-learn (v.0.22.1). The first two dimensions were used for visualization.

### Multi-omics factor analysis

Multi-omics decomposition by factor analysis was performed using the mofapy2 Python module (v0.5.6) (Argelaguet et al., 2018, 2020). Datasets with continuous measurements were selected for this analysis, i.e., drug responses, methylation, proteomics, and transcriptomics. For proteomic measurements, we used both the ProCan-DepMapSanger dataset and an independently acquired dataset (Nusinow et al., 2020) measuring an overlapping set of 290 cancer cell lines. Considering the strong separation of hematopoietic and lymphoid cell lines from the rest of the cell lines (***Figure 2a***), these were treated as a separate group in the analysis. Different numbers of factors were tested and *n* = 15 was chosen as it represented a trade-off between the total variance explained and the correlation between factors. Higher numbers of factors increased the correlation between factors and only marginally increased the variance explained, indicating that some factors were unnecessary. For the ProCan-DepMapSanger dataset, the mean sample intensity was regressed out prior to the factor analysis, thereby avoiding it being captured by any factor and artifactually increasing the total variance explained. Mofapy2 was run with convergence mode set to ‘slow’. Scale views and groups were set to ‘True’ to have a unit variance.

### Pairwise protein-protein correlations

We considered proteins with corresponding data also measured in the transcriptomics and CRISPR-Cas9 datasets (*n* = 6,347). For all pairwise protein combinations, we calculated Pearson’s *r* correlations between their protein, gene expression and essentiality measurements. A minimum of 15 complete observations was required to calculate the correlation, yielding 16,580,952 pairwise combinations. Protein-protein correlations were annotated using multiple sources of protein interactions: Comprehensive Resource of Mammalian Protein Complexes (CORUM) (Ruepp et al., 2010); Search Tool for the Retrieval of Interacting Genes/Proteins (STRING) (Szklarczyk et al., 2017); Biological General Repository for Interaction Datasets (BioGRID) (Chatr-Aryamontri et al., 2015); and Human Protein Interactome (HuRI) (Luck et al., 2020). For BioGRID, only physical interactions between proteins from humans were considered. For STRING, the most stringent threshold of the confidence score was chosen, and only interactions with a score ≥ 900 were considered. The average path length of the STRING PPI networks was 3.9. Protein-protein Pearson’s correlations were then used to estimate the capacity to recover interactions from the different resources by ranking in ascending order all correlations according to their *p*-value (x-axis) and drawing the cumulative distribution curve of the interactions found in the resource (y-axis). The area under the recall curve (AUC) was estimated using the corresponding function from the Python package scikit-learn (v0.24.2).

### DeeProM (DeepProteomicMarker)

We developed a multistep computational workflow, Deep Proteomic Marker (DeeProM), to identify protein biomarkers of cancer vulnerabilities. The analysis steps in DeeProM are fourfold. First, it prioritizes drug responses and CRISPR-Cas9 gene essentialities that can be confidently predicted using proteomic profiles. Second, it prioritizes strong drug responses and gene essentialities that are specific to small subsets of cancer cell lines. Third, it prioritizes protein biomarkers that show significant associations with drug responses or gene essentialities. Fourth, it prioritizes protein biomarkers that are present in specific tissues. The source code is provided in the GitHub repository (see **Data and Software Availability**).

#### DeepOmicNet

DeeProM is powered by a deep neural network architecture, DeepOmicNet, to predict drug responses and CRISPR-Cas9 gene essentialities. DeepOmicNet ranks drugs and gene essentialities based on predicted cellular responses using the ProCan-DepMapSanger dataset as the input. Both the proteomic and the drug response datasets contain missing values, while the gene essentiality dataset provides a complete data matrix. DeepOmicNet models these missing values accordingly. Multilayer perceptron (MLP) is a classic neural network architecture that has been used by default for deep learning in numerous biomedical studies (Zhang et al., 2019). To enhance the predictive performance of MLP with proteomic data, we modified its network architecture and developed DeepOmicNet with the following three major improvements:

#### Grouped bottleneck

DeepOmicNet uses grouped bottlenecks to avoid fully connected layers, which involves a large number of parameters being optimized. Compared with a fully-connected layer, breaking the connections into smaller groups allows the network to be more memory efficient, thus enabling wider or deeper layers. A weight matrix *W* ∈ *R*^*(k×k)*^ containing *k*^*2*^ parameters with grouped bottlenecks can not only reduce the number of parameters, but also provide better predictive performance. Instead of connecting all pairs of neurons, neurons can be broken into groups, and only neurons within the same group are connected between layers (***Figure S3b***). The group size *g* can be set as any number that is divisible by the hidden layer width *k*. When *g=k*, all neurons are treated as one group, which reduces to a normal fully-connected layer. Multiple configurations were tested and the optimal group size *g* was set to 2. The number of parameters for one layer with grouped bottlenecks is significantly reduced from *k*^*2*^ to *k/2×2*^*2*^*=2k*. The number of parameters with grouped bottlenecks is calculated as *k/g×g*^*2*^*=g×k*, thus the run-time complexity is decreased from quadratic to linear.

#### Skip connections

The complete network architecture is visualized in ***Figure S3b***. Neurons between every two consecutive layers are connected in MLP, which is computationally intensive and suboptimal for model training. To mitigate this problem, DeepOmicNet utilizes skip connections (He et al., 2016) to connect alternate layers. Let *x*∈*R*^*k*^ be the vector of the *i*^*th*^ hidden layer of a real coordinate space of dimension *k* (corresponds to the number of neurons in a layer, also known as the layer width), the value of *x*_*i*_ is calculated with skip connections as:

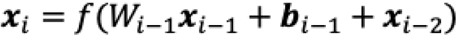

where *f* is the activation function, *W*_*i*−1_∈*R*^(*kxk*)^ is the weight matrix, *b*_*i*−1_∈*R*^*k*^ is the bias vector and *x*_*i*−2_∈*R*^*k*^ is the hidden layer ahead of the hidden layer *x*_*i*−1_. The value of the hidden layer *i-2* is fed into the hidden layer *i* by skipping the hidden layer *i-1*, resulting in skip connections (***Figure S3b***). Each hidden layer is set with the same width *k*, which is a hyperparameter for model tuning, and usually is chosen to be slightly smaller than the input feature dimension. In DeepOmicNet, a sigmoid function was chosen to be the activation function *f*, because it outperformed the rectified linear activation function and the hyperbolic tangent function.

#### Loss function

DeepOmicNet is trained with mini-batches using a customized mean squared error (MSE) as the loss function. DeepOmicNet is applied to predict both drug responses and CRISPR-Cas9 gene essentialities. For the cell line *m*, the loss for the target variable *n* is defined as:

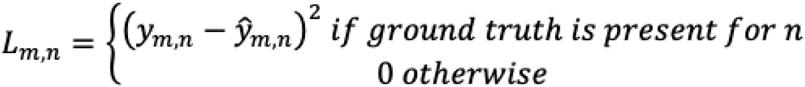

where *y*_*m,n*_ is the ground truth label of the target *n* (either drug response or gene essentiality) and *ŷ*_*m,n*_ is the predicted value of the target *n*. The label *y*_*m,n*_ is missing if a particular cell line *m* was not screened with drug *n*.

In addition to the three major improvements, other characteristics of DeepOmicNet include the following:

#### Missing values

One specific challenge of DIA-MS based proteomics is the missing values in the data matrix (Poulos et al., 2020; Webb-Robertson et al., 2015). Imputation is widely used but often leads to distortion to some extent (Wei et al., 2018). For DeepOmicNet, missing values were replaced by zeros, thus allowing the neural network to ignore the weight update for these inputs.

#### Hyperparameter tuning

Hyperparameters including model width, depth, learning rate and batch size were tuned to achieve the highest predictive performance. Pearson’s *r* between true and predicted values is used as the evaluation metric. Hyperparameters that resulted in the highest performance in five-fold cross-validation of the 80% training data were chosen for the final evaluation on the 20% independent test set. The chosen hyperparameters can be found in the configuration files in the source code (see **Data and Software Availability**).

#### Thresholding

The thresholds are set to Pearson’s *r* > 0.4 and Pearson’s *r* > 0.3 to prioritize highly predictive drug responses and CRISPR-Cas9 gene essentialities, respectively. This yielded 67 drug responses and 62 gene essentialities.

#### Identification of drug responses and CRISPR-Cas9 gene essentialities

DeeProM prioritizes drug responses and CRISPR-Cas9 gene essentialities that are likely to be non-toxic to normal cells. Since the cell lines used in this study were derived from cancer (the majority) or viral transformation, we approximated this task by finding drug responses and gene essentialities that were selective for only a small fraction of the cell lines, which indicates that the drug response or gene essentiality is less likely to be toxic to normal cells. Ranking of strongly selective drug responses and gene essentialities was performed using Python package scipy (v1.5.2) skew function, which calculates the Fisher-Pearson’s coefficient of skewness. Skewness values of -1 and -2 were used for drug responses and gene essentialities, respectively.

#### Linear regression models for protein feature identification

Associations between protein and phenotypic measurements, drug responses and gene essentialities were performed using linear regression models (sklearn v0.24.2 class LinearRegression). Several technical and biological covariates were added to the model to remove potentially spurious associations. First, we built the following technical covariates into the model: (i) the growth rate of the cell lines; (ii) cell culture medium, D/F12 (DMEM/F12: 10% FBS, 1% PenStrep) or R (RPMI1640: 10% FBS, 1% PenStrep, 4.5 mg/ml Glucose, 1 mM Sodium Pyruvate); (iii) cell line growth properties, i.e., adherent, semi-adherent or suspension; (iv) sample mean protein replicates Pearson’s correlation; (v) for the CRISPR-Cas9 gene essentiality only, we considered the institute of origin of the CRISPR-Cas9 screen, i.e., Wellcome Sanger or Broad Institute; and (vi) for the drug response models only, we considered the cell line mean IC_50_ across all drugs. Discrete covariates were represented as dummy binary variables. Second, to identify associations that were exclusively found at the protein level, we added the following gene expression covariates to the model: (i) the first ten gene expression principal components using the Python package scikit-learn (v0.24.2); and (ii) the corresponding transcript level of the protein being tested. Formally, we fit the following linear regression model for each drug response/gene essentiality–protein pair:

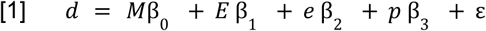

where, *d* represents a *n* x 1 vector of the drug response IC_50_ (*n* = 710 drugs) for 947 cell lines or CRISPR-Cas9 gene essentiality log_2_ fold changes (*n* = 17,486 genes) for 534 cell lines; *M* is the *n* x *k* matrix of covariates (k = 11 covariates); *E* is a *n* x *m* matrix containing the first (*m* = 10) principal components of the gene expression dataset; *e* is a vector of size *n* x 1 containing the transcriptomics measurements of the corresponding protein *p*; *p* is a vector of size *n* x 1 containing the protein measurements; and ε is the error vector of size *n* x 1. For each protein, cell lines with missing values were dropped from the modeling. For drug response, missing values were replaced by the drug mean IC_50_. The number of cell lines with complete information per fit was provided. The model was fitted by minimizing the residual sum of squares to estimate the parameters β_*n*_ of each variable. In total, there were 710 × 6,692 = 4,751,320 drug-protein pairs and 17,486 × 6,692 = 117,016,312 possible CRISPR-Cas9 gene essentiality-protein pairs, however, both drug response and CRISPR-Cas9 data covered various subsets of cell lines. We required a minimum number of 60 cell lines to test the association. As a result, a total of 4,218,788 drug-protein and 86,584,537 CRISPR-Cas9-protein tests were performed.

Statistical assessment of the improvement of adding protein measurements to the linear regression was performed using a likelihood ratio test between the full model [Equation 1] and the null model, which excludes the protein measurement and its parameter β_2_. Likelihood ratio test’s *p*-value was estimated using a chi-square distribution with one degree of freedom. Adjustment for multiple testing was performed per drug or CRISPR-Cas9 gene essentiality using the Benjamini-Hochberg procedure to control the false discovery rate (FDR). Associations with FDR < 0.1 for models with covariates, or FDR < 0.001 for models that do not use covariates, were considered as significant associations.

#### Tissue type level filtering

To investigate drug-protein and CRISPR-Cas9-protein associations for a given tissue type, we used metrics derived from DeepOmicNet, Fisher Pearson’s coefficient of skewness and linear regression to prioritize drug responses and CRISPR-Cas9 gene essentialities according to the thresholds described above. The overlap of these three methods was used as the final result, which included 7,698 drug-protein and 5,823 CRISPR-Cas9-protein associations. Finally, we applied additional filters to further prioritize associations that are unique to certain tissue types, yielding the final list of 108 drug-protein associations for 18 drugs and 1,538 CRISPR-Cas9-protein associations for 38 genes (***Figure 4c***). The filtering steps are described below:

##### Step 1

Tissue types with < 20 protein measurements were filtered out.

##### Step 2

For each significant association identified from 7,698 drug-protein and 5,823 CRISPR-Cas9-protein associations, Pearson’s *r* was calculated. Using protein data for each tissue type, the significance level was set to 0.1 and Pearson’s *r* was defined as *r*_*(target, protein)*_^*tissue*^ for a target-protein association, where protein indicates a protein of interest, target indicates either a drug response or CRISPR-Cas9 gene essentiality. We then calculated the Pearson’s *r* for the gene that encodes each protein, defined as *r*_*(target, RNA)*_^*tissue*^. To prioritize associations that were uniquely identified at the protein level, the difference *d* for a given tissue type between target-protein and target-RNA associations was set to be larger than 0.15, where *d* = |*r*_*(target, protein)*_^*tissue*^ |*-*|*r*_*(target, RNA)*_^*tissue*^ |. This prioritizes associations that have either strong positive or strong negative correlations at the protein level but have weak correlations around zero at the RNA level. Rare cases where *r*_*(target, protein)*_^*tissue*^ and *r*_*(target, RNA)*_^*tissue*^ have opposite signs and *d* is close to 0 were not considered.

##### Step 3

For drug-protein associations, a large value of *d* alone is insufficient to select candidate associations, because a drug may be entirely ineffective for all the cell lines in a particular tissue type. Therefore, we applied an additional filter to ensure that a drug is effective on the cell lines for which the protein abundance is high. That is, for a drug-protein association to be included for a given tissue type, the median IC_50_ of the 20% of cell lines with the highest corresponding protein abundance must be lower than the maximal concentration for that drug.

##### Step 4

The remaining associations were ranked in descending order according to *d*.

### Comparison between DeepOmicNet and traditional machine learning models on drug response

DeepOmicNet was compared against traditional machine learning models, including elastic net and Random Forest. A total of 947 cell lines were randomly separated into a training set comprising 80% of the cell lines, and a test set with the remaining cell lines for unbiased evaluation. Grid search was used to find the best hyperparameters for elastic net and Random Forest in the training set by cross-validation. Hyperparameter tuning for DeepOmicNet was performed manually due to the limit of graphics processing unit (GPU) memory. For each model, missing values were imputed using the method that gave the best prediction based on cross-validation. Specifically, four imputation methods were considered, including imputation by minimum, first percentile of the whole input matrix, mean and zero. Based on the predictive performance of models in cross-validation, imputation by one percentile of the whole matrix was chosen for the elastic net and Random Forest, and imputation by zero was used for DeepOmicNet. This strategy yielded the best prediction accuracy in comparison with other imputation and normalization methods, such as imputation with k-nearest-neighbor, mean values of proteins and zeros. A cut-off was set at a minimum of 300 screened cell lines for the drug response dataset to filter out drugs without sufficient data. A simplified version of DeepOmicNet without grouped bottlenecks was used for omics data other than proteomic data due to the large input dimension. The Python package scikit-learn (v.0.22.1) was used to train elastic net and Random Forest models. DeepOmicNet was implemented and trained using PyTorch (v.1.4.0).

### Machine learning for CRISPR-Cas9 lethality prediction

Due to the limit of GPU memory, elastic net, Random Forest and DeepOmicNet were applied only to transcriptomic and proteomic data to predict CRISPR-Cas9 gene essentialities. The same computational strategy for drug response prediction was used to predict CRISPR-Cas9 gene essentialities.

### Predictive power comparison with the Cancer Cell Line Encyclopedia (CCLE)

The predictive power of machine learning models for two proteomic datasets (ProCan-DepMapSanger and CCLE (Nusinow et al., 2020)) were compared independently on three drug response datasets (Sanger, CTD2 and PRISM) and the CRISPR-Cas9 gene essentiality dataset (Behan et al., 2019; Meyers et al., 2017; Pacini et al., 2021). The analysis was performed using the 290 overlapping cell lines to ensure a fair comparison. The proteomic (Nusinow et al., 2020) and drug response (CTD2 and PRISM) (Corsello et al., 2020; Rees et al., 2016; Seashore-Ludlow et al., 2015) datasets were retrieved from the DepMap portal (https://depmap.org/portal/). DeepOmicNet was used to compare the predictive power of models for CRISPR-Cas9 gene essentialities, and Random Forest was used for drug response prediction due to the limited number of cell lines for certain drugs. AUC instead of IC_50_ was used for the drug response (PRISM) dataset due to a large proportion of drugs having no IC_50_ values provided.

### Random downsampling analysis of drug response prediction

For downsampling analysis, the full set of 8,498 proteins were randomly downsampled using a step decrease of 500 proteins (***Figure 7a***). Each step was repeated ten times and for each iteration, results from five-fold cross-validations and an unbiased test were included in evaluating predictive power. Therefore, each downsampling step used ten different random subsets of proteins for six distinct experiments (the five-fold cross-validation and one unbiased test). The predictive power of each DeepOmicNet model was evaluated for each protein set and each iteration, with confidence intervals summarizing the results across the ten iterations. This random downsampling procedure was also performed with step sizes of 250 proteins for proteins in Categories A, B and C (***Figure 7e***).

## Data and Code Availability

All data will be made available upon publication. The raw mass spectrometry proteomic data and accompanying files will be available in the ProteomeXchange Consortium via the PRIDE (Perez-Riverol et al., 2019) partner repository. Drug response measurements will be made public through www.cancerrxgene.org portal. The proteomics and drug response datasets will be integrated with existing cell line data at www.cellmodelpassports.sanger.ac.uk portal. Code related to the analyses will be made available on github upon publication.

## Supplemental Information

### Supplementary Tables

Refer to the accompanying Excel files.

***Table S1***. Annotated list of cancer cell lines and MS files used in this study.

***Table S2***. Protein measurements across 949 cancer cell lines.

***Table S3***. Data underlying cell type-enriched proteins and mutation analysis.

***Table S4***. MOFA analysis cell line factors and feature weights.

***Table S5***. Protein–protein correlations.

***Table S6***. Results of DeeProM before tissue type level filtering is applied.

***Table S7***. Results of DeeProM for tissue-specific biomarkers.

***Table S8***. Data underlying MS processing steps.

### Supplementary Figures

**Figure S1.**
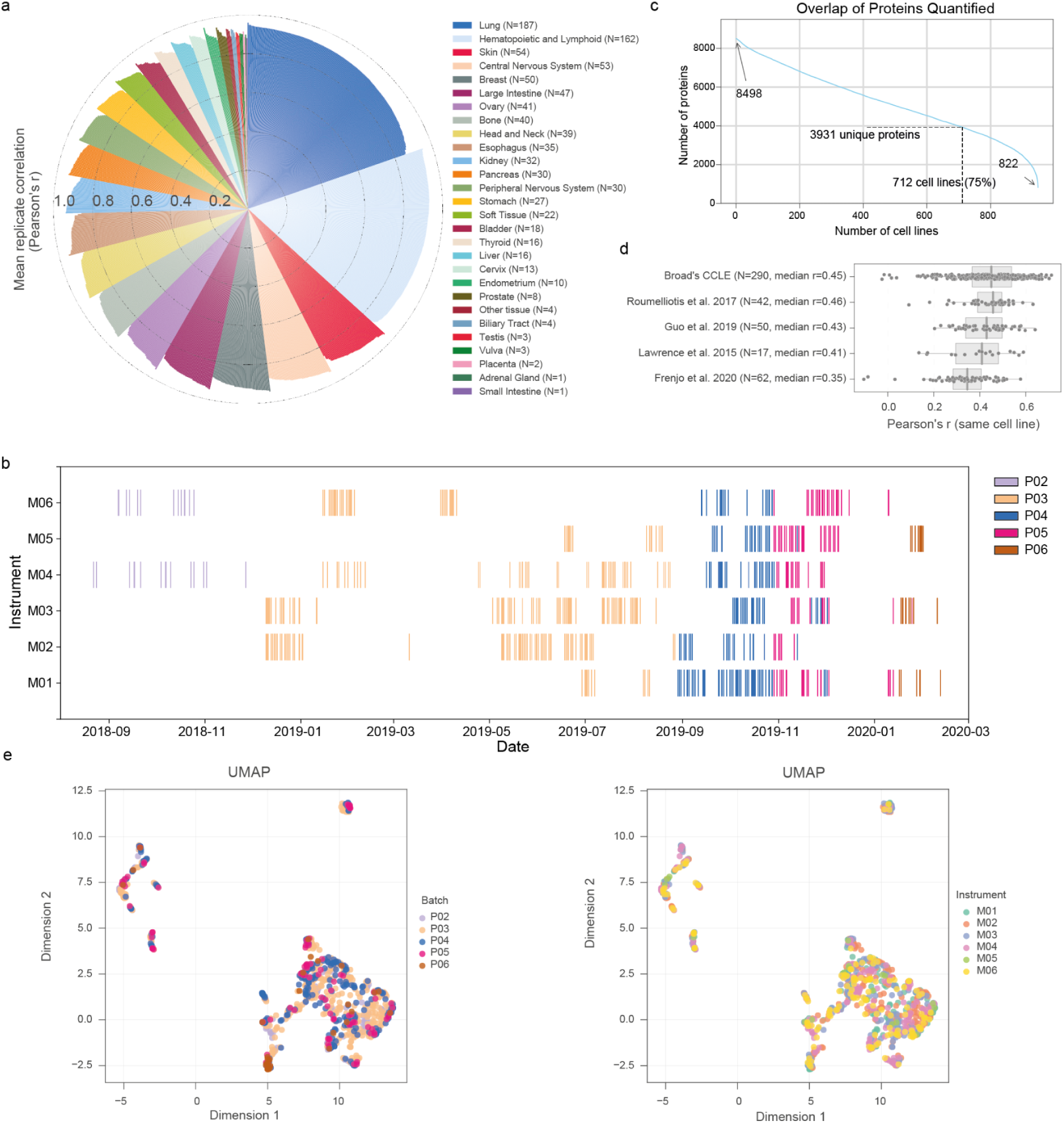
A pan-cancer proteomic map of 949 human cancer cell lines by DIA-MS. **a**, Mean Pearson’s r for replicates of each cancer cell line, colored by tissue of origin. **b**, Timeline of MS data acquisition across mass spectrometers, colored according to processing batches (P02 - P06). **c**, Frequency of proteins identified across the 949 cancer cell lines. **d**, Correlation by Pearson’s r of ProCan-DepMapSanger dataset against independent proteomic datasets that comprise subsets of the same cell lines. Box-and-whisker plots show 1.5 x interquartile ranges and centers indicate medians. **e**, UMAP dimensionality reduction of cell line proteomes colored by processing batches (left) and mass spectrometer (right).

**Figure S2.**
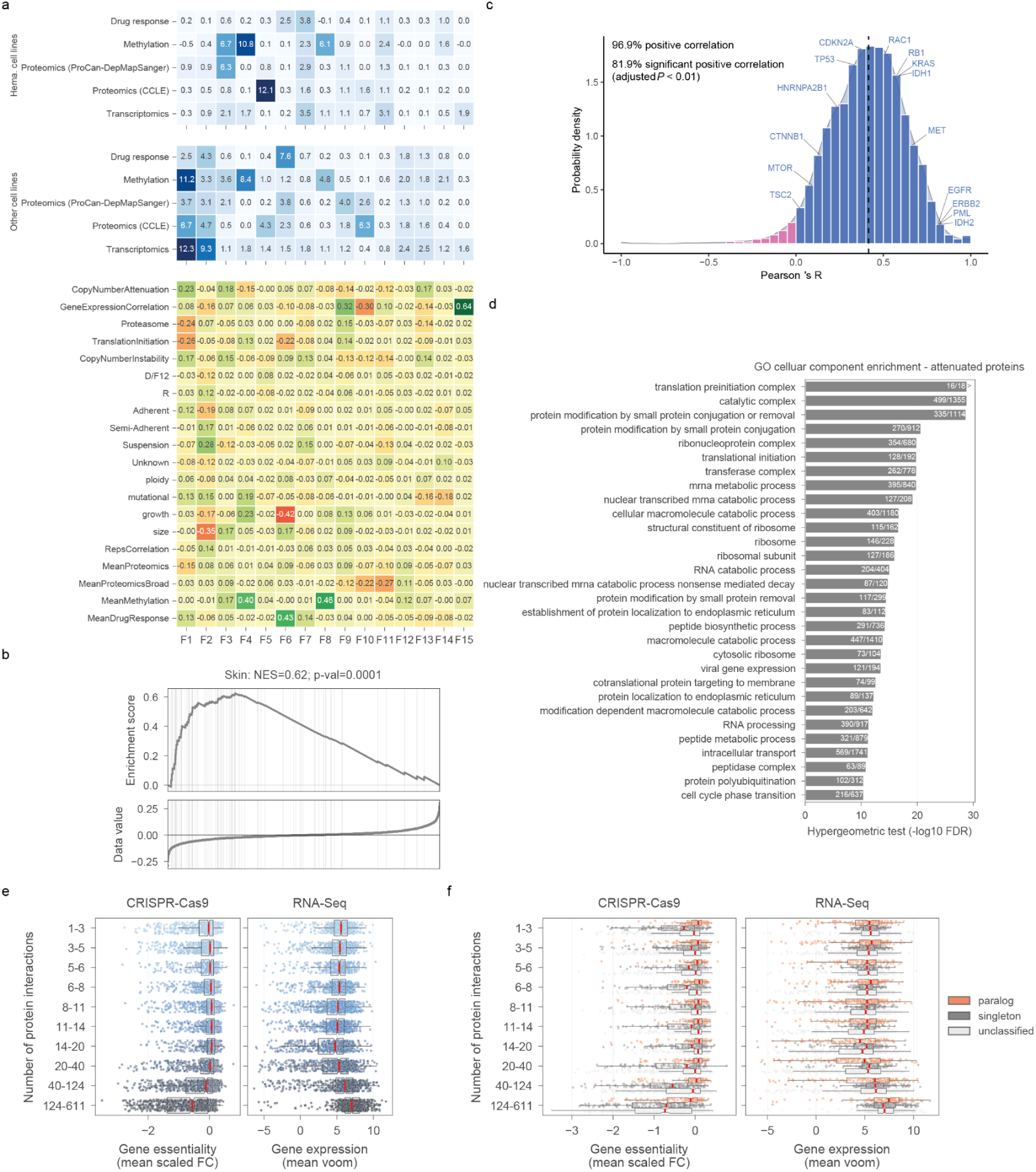
MOFA and post-transcriptional regulation. **a**, Similar to **Figure 3a**, MOFA factors across molecular and phenotypic cancer cell line datasets, including ProCan-DepMapSanger. Hematopoietic and lymphoid cells are grouped and trained separately from the other cell lines corresponding to each factor (column). The upper two heatmaps (blue) report the portion of variance explained by each factor (columns) in each dataset. The lower heatmap reports Pearson’s r between each learned factor and various molecular characteristics of the cancer cell lines. **b**, GSEA demonstrating enrichment of skin cell type-enriched proteins in MOFA Factor 12. **c**, Per-gene Pearson’s r between protein and RNA expression for all proteins quantified. Mean correlation (r = 0.42) is indicated by a dashed line. The locations of several cancer-related genes are shown. **d**, Enrichment analysis of proteins that were highly attenuated (n = 1,215), as determined by correlations between protein and copy number, and gene expression and copy number, in terms of Pearson’s r. The top most significantly enriched sets are annotated. **e, f**, Proteins were grouped by their number of significant positive correlations by Pearson’s r for putative protein interactions (FDR < 5%, r > 0.5). For each protein, the respective mean scaled CRISPR-Cas9 gene essentiality fold-change (FC) and gene expression (RNA-seq voom) measurement are calculated. **e**, The distribution across all proteins is represented and **f**, proteins are subgrouped into paralog, singleton and unclassified, as previously determined (Dandage and Landry, 2019). Box-and-whisker plots show 1.5 x interquartile ranges and centers indicate medians.

**Figure S3.**
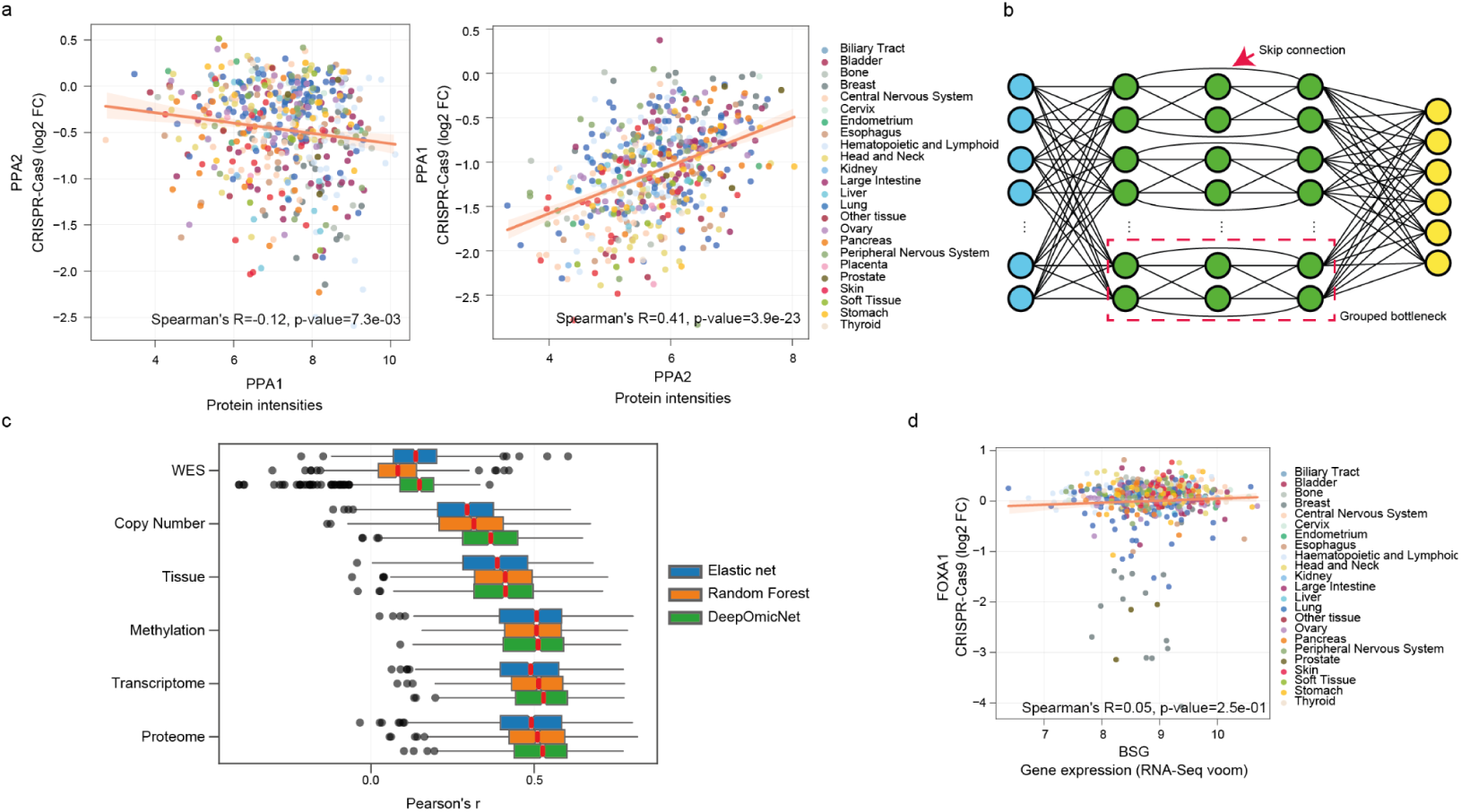
Drug-protein and CRISPR-Cas9-protein associations and DeeProM analysis pipeline. **a**, Synthetic lethal association between PPA2 and PPA1. Left panel, scatter plot between protein intensities of PPA1 and PPA2 CRISPR-Cas9 gene essentiality scores. Right panel, scatter plot between protein intensities of PPA2 and CRISPR-Cas9 gene essentiality scores of PPA1. Cell lines are colored by tissue types. **b**, Neural network architecture of DeepOmicNet. In addition to the basic MLP architecture, skip connections with grouped bottlenecks were added to provide a deeper and wider network. Blue circles represent input neurons, green represents hidden layer neurons and yellow represents output neurons. **c**, Comparison of observed drug responses and predicted drug responses by DeeProM, elastic net and Random Forest. WES, mutation data from whole exome sequencing; Copy number, copy number profiles; Tissue, categorical variable representing the cell line’s tissue of origin; Methylation, promoter region methylation level; Transcriptome, RNA-seq data; Proteome, the ProCan-DepMapSanger dataset. Box-and-whisker plots show 1.5 x interquartile ranges and centers indicate medians. **d**, Scatter plot between FOXA1 CRISPR-Cas9 gene essentiality scores and BSG gene expression measurements. Data points are colored according to tissue type.

**Figure S4.**
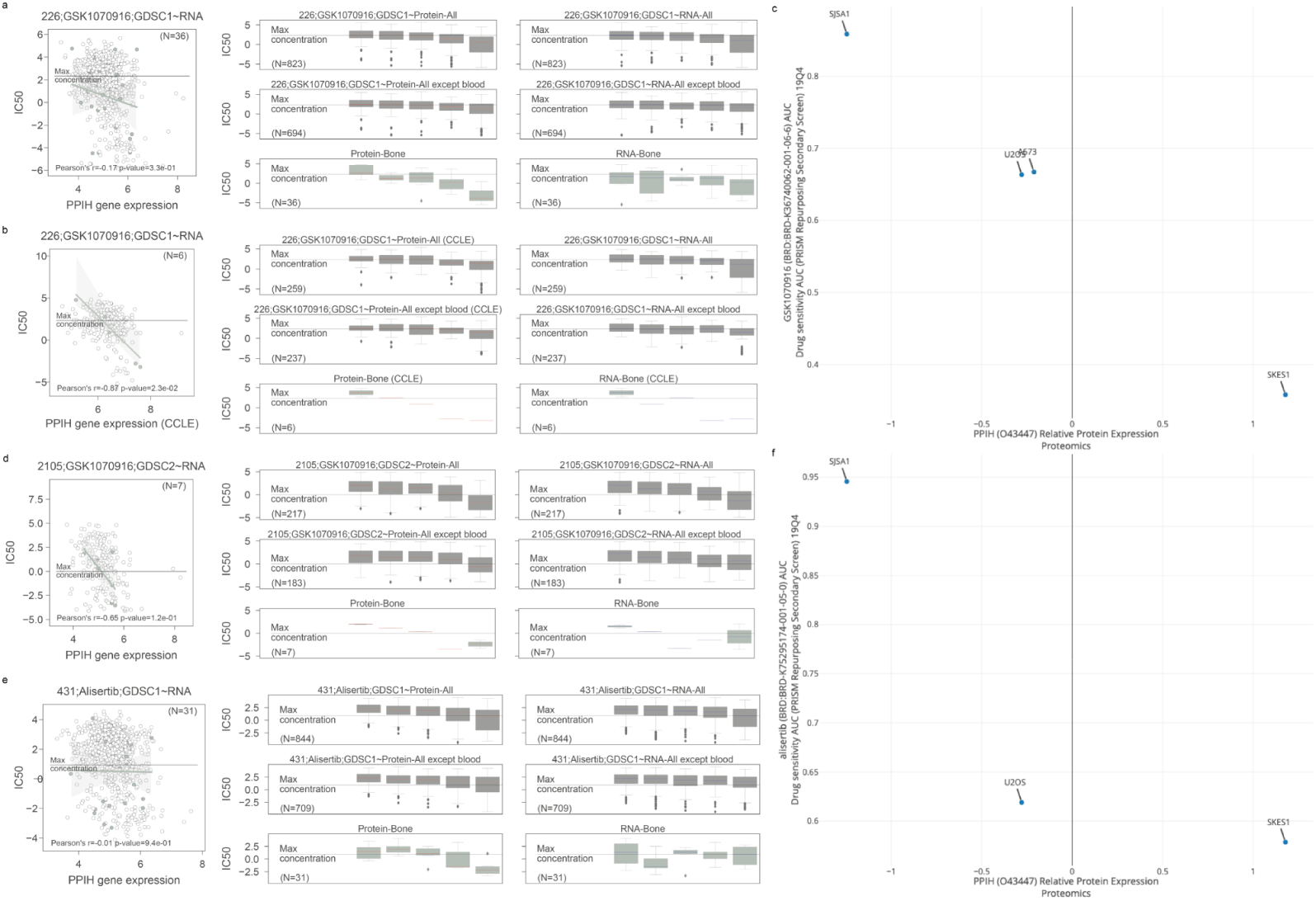
Tissue-level protein biomarkers for GSK1070916 and Alisertib. **a, b, d, e**, Scatter plots on the left of each panel show the relationship between the drug and the underlying RNA expression of the protein biomarker in the tissue type indicated (green; all other cell lines are shown in gray). The number of cell lines and Pearson’s r from the highlighted tissue type are annotated at the top right and bottom left corners, respectively. Box-and-whisker plots summarize the information from the scatter plots in **Figure 5d-g** (protein; left) and **Figure S4a,b,d,e** (RNA; right). The protein intensity is divided into five bins from low to high, and the corresponding IC_50_ values in the natural logarithmic scale are shown for each quartile. The first row of plots shows the relationship for all cell lines, and non-hematopoietic cell lines are shown in the second row. The highlighted tissue type is shown in the third row. Box-and-whisker plots show 1.5 x interquartile ranges and centers indicate medians. **a**, The association between PPIH and GSK1070916 drug response from GDSC data in cell lines from bone. **b**, The same as **a**, but instead using CCLE gene expression data. **d**, The same as **a**, but instead using GDSC2 drug response data. **e**, The same as **a**, but instead showing association between PPIH and Alisertib drug response (from GDSC data). **c, f**, The association between PPIH protein abundance and response to **c**, GSK1070916 and **f**, Alisertib, using CCLE proteomic dataset and PRISM drug response data, with figures obtained from the DepMap Portal.

**Figure S5.**
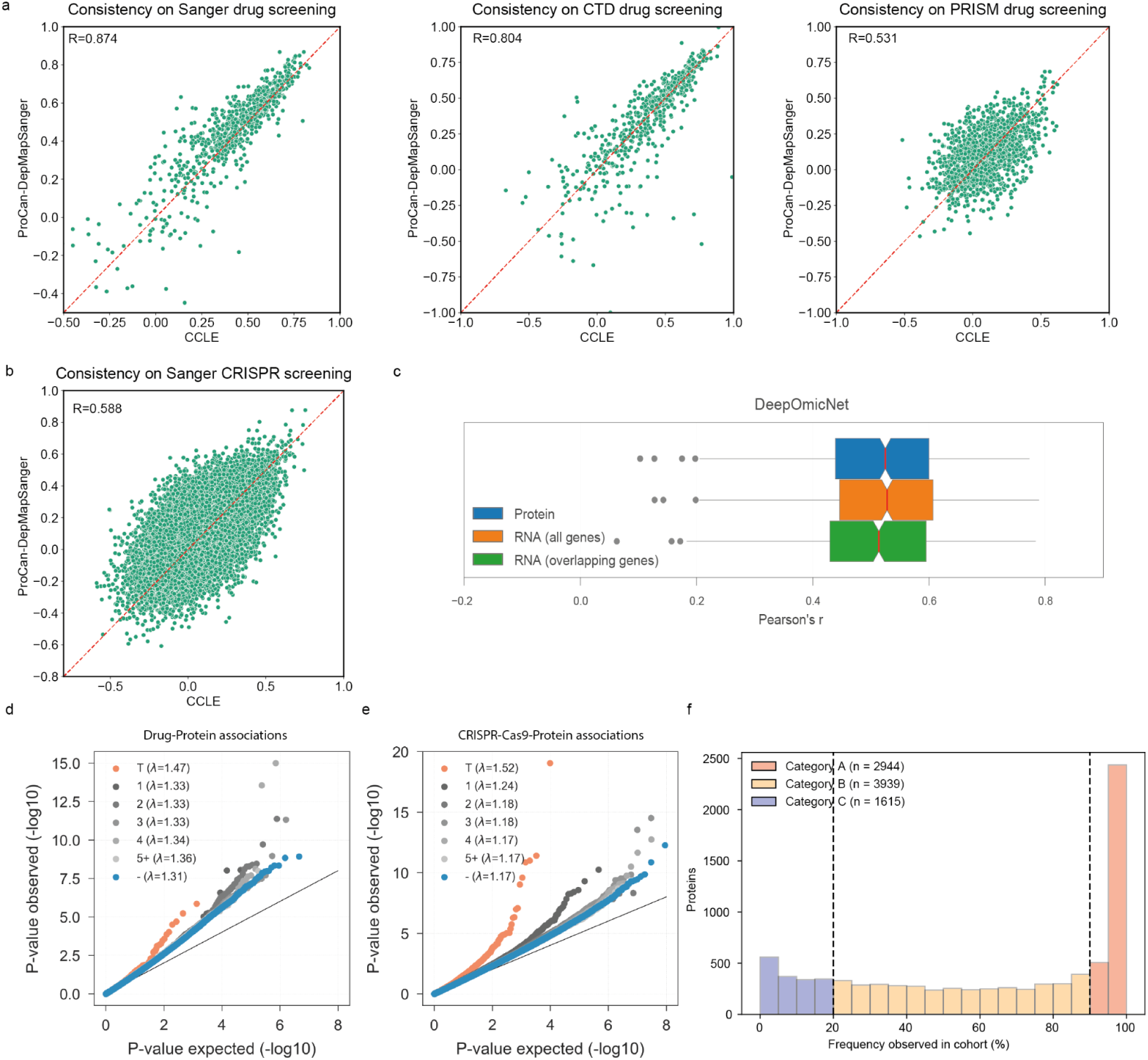
Predictive power benchmarks and comparisons. **a**, Comparison of DeepOmicNet mean predictive power across three independent drug response datasets, trained using the ProCan-DepMapSanger and the CCLE proteomic dataset. **b**, Similar to **a**, models trained to predict CRISPR-Cas9 gene essentiality profiles. **c**, Comparison of the predictive power of the DeepOmicNet model trained using the ProCan-DepMapSanger dataset and RNA expression data, both for all genes or for only genes that have their corresponding proteins quantified (overlapping genes). **d**, Drug responses (drug-protein) and **e**, CRISPR-Cas9 gene essentiality (CRISPR-Cas9-protein) associations, identified with linear regression models without taking gene expression as covariates. See **Figure 7c-d** for a general description of the plot. **f**, Histogram showing the distribution of the frequency of proteins detected in each of the cell lines in the cohort, with Category A, B and C proteins indicated. The dashed vertical lines indicate the frequency thresholds for defining the categories.

